# Single-cell transcriptomic profiling of healthy and fibrotic adult zebrafish liver reveals conserved cell identities and stellate cell activation phenotypes with human liver

**DOI:** 10.1101/2021.08.06.455422

**Authors:** Joshua K. Morrison, Charles DeRossi, Isaac L. Alter, Shikha Nayar, Mamta Giri, Chi Zhang, Judy H. Cho, Jaime Chu

## Abstract

Liver fibrosis is the excessive accumulation of extracellular matrix that can progress to cirrhosis and failure if untreated (1). The mechanisms of fibrogenesis are multi-faceted and remain elusive with no approved antifibrotic treatments available (2). Here we use single-cell RNA sequencing (scRNA-seq) of the adult zebrafish liver to study the molecular and cellular dynamics of the liver at a single-cell level and demonstrate the value of the adult zebrafish as a model for studying liver fibrosis. scRNA-seq reveals transcriptionally unique populations of hepatic cell types that comprise the zebrafish liver. Joint clustering with human liver scRNA-seq data demonstrates high conservation of transcriptional profiles and human marker genes in zebrafish cell types. Human and zebrafish hepatic stellate cells (HSCs), the driver cell in liver fibrosis (3), specifically show conservation of transcriptional profiles and we uncover Colec11 as a novel, conserved marker for zebrafish HSCs. To demonstrate the power of scRNA-seq to study liver fibrosis, we performed scRNA-seq on our zebrafish model of a pediatric liver disease with characteristic early, progressive liver fibrosis caused by mutation in mannose phosphate isomerase (MPI) (4–6). Comparison of differentially expressed genes from human and zebrafish *MPI* mutant HSC datasets demonstrated similar activation of fibrosis signaling pathways and upstream regulators. CellPhoneDB analysis revealed important receptor-ligand interactions within normal and fibrotic states. This study establishes the first scRNA-seq atlas of the adult zebrafish liver, highlights the high degree of similarity to the human liver, and strengthens its value as a model to study liver fibrosis.

**Significance Statement:** To our knowledge, this is the first single-cell characterization of the adult zebrafish liver, both in a normal physiologic state and in the setting of liver fibrosis. We identify transcriptionally distinct zebrafish liver cell populations and a high degree of transcriptional conservation between human and zebrafish cells across the majority of hepatic cell types. Furthermore, using this scRNA transcriptome, we identify key signaling pathways in zebrafish HSCs that are replicated in human HSCs and implicated in the regulation of liver fibrosis. Our work provides a useful resource that can be used to aid research using the zebrafish liver and asserts the usefulness of the adult zebrafish to study liver fibrosis.

## Introduction

The liver is an essential organ for regulating multiple metabolic and homeostatic processes; these functions are disrupted in liver injury and disease resulting in high morbidity and mortality (7, 8). Liver fibrosis is the excessive accumulation of extracellular matrix (ECM) that is produced in response to chronic liver injury which can progress to cirrhosis and liver failure if untreated (1).Despite the increasing prevalence and high morbidity of progressive liver fibrosis, there are currently no approved antifibrotic treatments (2). Utilizing animal models to study liver function and disease is essential to better understand liver cell biology, identify new therapeutic targets, and establish pre-clinical animal models to test potential therapies.

The zebrafish has increasingly become accepted as a powerful model to study liver pathologies, including steatosis, fibrosis, and hepatocellular carcinoma (9–11). The zebrafish liver is remarkably similar to the human liver, with conserved cellular composition and functionality. Furthermore, the zebrafish liver is fully functional by 5 days post-fertilization, making it a fast, high-throughput tool for disease modeling (9, 10). The majority of studies have focused on larval zebrafish to model acute liver injury (12–14), but adult zebrafish have been underutilized to study chronic liver diseases such as fibrosis.

Mannose phosphate isomerase (MPI) is an essential enzyme that interconverts mannose 6-phosphate and fructose 6-phosphate (Man6P ←→ Fru6P). MPI mutation in humans result in a congenital disorder of glycosylation (MPI-CDG) that is characterized by a congenital form of liver fibrosis (4–6). We previously generated a zebrafish mutant *mpi* line (*mpi*^*mss7*^) that recapitulated key features of MPI-CDG, including characteristic liver fibrosis (15). We found that MPI depletion could directly activate hepatic stellate cells (HSCs), the main driver cell of liver fibrosis, but the effects of MPI loss in other liver cell types and the impact of these effects on the regulation of liver fibrosis is not known (16).

Single-cell RNA sequencing (scRNA-seq) has emerged as a powerful tool to identify key marker genes and pathways in various cell types, uncover unique cell populations and subpopulations, and allows for the study of rare cell populations in tissues (17, 18). Recent high-impact studies on human livers have utilized scRNA-seq to reveal the existence and investigate the functions of unique subpopulations of various hepatic cell types including macrophages, epithelial progenitor cells, and myofibroblasts (19–23). In contrast, there has been no reported scRNA-seq data on the adult zebrafish liver, including fibrotic liver disease, which limits the ability to study molecular and cellular dynamics in the mature liver at a single-cell level.

Here we present the first full characterization of the adult zebrafish liver transcriptome at the single-cell level and demonstrate its utility through comparison with the *mpi*^*+/mss7*^ zebrafish model of fibrosis to uncover potential mechanisms of fibrogenesis. We further compare and contrast the transcriptional cell identities with adult human liver, specifically zebrafish HSC signatures to that of human HSCs in physiologic conditions, as well as in fibrosis using MPI-depleted conditions. This adult zebrafish liver cell atlas can be a valuable resource to expand the use of adult zebrafish as a tool to study liver fibrosis.

## Results

### Creation of a single cell atlas of the adult zebrafish liver

To develop a single-cell atlas of adult zebrafish liver, livers were dissected from 18-month-old wild-type zebrafish (WT; N = 3). Single-cell suspensions were generated following previously published protocols (24), with cell viabilities from each sample ranging from 72 to 81%. After mitochondrial RNA percentage, unique molecular identifier (UMI), and gene cutoffs were implemented, a range of 3,742 to 5,544 profiled cells (N = 13,630 cells) from each sample were used for further analysis. Samples were integrated to perform joint clustering using the standard Seurat pipeline (25–28). Nineteen transcriptionally distinct clusters were identified (**Fig. 1A-B, Table S1**), and each cluster contained cells from all 3 WT samples, supporting that these clusters are representative of true biological populations (**Fig. S1A-B, Table S2**).

**Figure 1.**
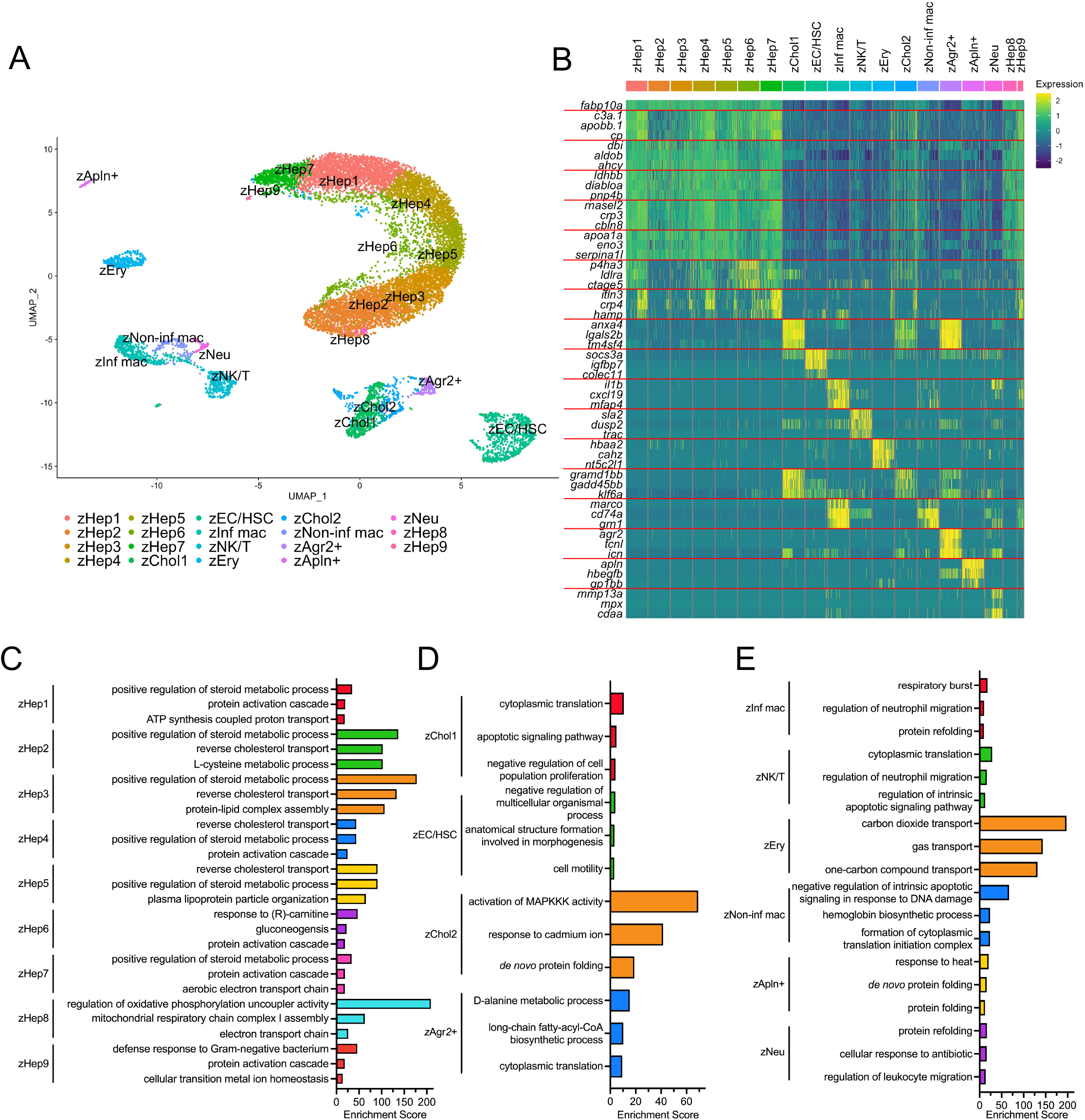
Single-cell transcriptome of the adult zebrafish liver. **A**. scRNA-seq of WT adult zebrafish liver cells (N = 3). Uniform manifold approximation and projection (UMAP) shows 19 unique clusters. **B**. Heatmap of gene expression for three of the top differentially expressed genes in each cluster. **C-E**. Bar graph of the top three GO processes enriched for in each cluster of the WT zebrafish adult liver atlas. **Abbreviations**: zHep – hepatocyte, zChol – cholangiocyte, zEC/HSC – endothelial cell/hepatic stellate cell, zInf mac – inflammatory macrophage, zNK/T – NK cell/T cell, zEry – erythrocyte, zNon-inf mac – non-inflammatory macrophage, zAgr2+ – *agr2+* cell, zApln+ – *apln+* cell, zNeu – neutrophil.

Hepatocytes were the most abundant cell type recovered from zebrafish liver (73% of all liver cells captured), identified by high expression of *fabp10a*, an established marker that is restricted to hepatocytes in the zebrafish liver (**Fig. 1B**) (29, 30). Other hepatocyte-specific marker genes, *tfa* (31) and *cp* (32), were detected exclusively in hepatocytes and confirmed the ability to discriminate hepatocytes from other cell populations in our zebrafish liver dataset (**Fig. S1C-D**). Furthermore, based on marker gene expression, hepatocytes could be further separated into distinct clusters, comprising 9 of the 19 total clusters identified (zHep1-9) (**Fig. 1A-B, Table S1**). Gene ontology (GO) enrichment analysis revealed that these hepatocyte clusters were generally involved with metabolic pathways, including steroid metabolic processes and cholesterol transport (**Fig. 1C**). Interestingly, clusters zHep1, zHep7, and zHep8 were enriched for pathways involved in oxidative phosphorylation, while clusters zHep2, zHep3, zHep4, and zHep5 were enriched for pathways involved in cholesterol, steroid, and lipid metabolism and transport. Cluster zHep6 was enriched for processes involved in fatty acid transport and glucose metabolism. Cluster zHep9 was enriched for immune processes (**Fig. 1C**), but after further analysis of the number of UMIs detected and marker gene expression (**Fig. 1B, S1E**) these cells appeared to be heterogenous multiplets of hepatocytes and immune cells. Collectively, these data demonstrate that the adult zebrafish liver is comprised of multiple unique hepatocyte populations with distinct functions, similar to the specialization of human hepatocytes (19, 33, 34).

Cholangiocytes were the second most abundant cell type detected, encompassing roughly 8% of all liver cells captured and making up 3 out of the 19 total clusters (zChol1, zChol2, and zAgr2+) (**Fig. 1A-B**). Cholangiocyte clusters were identified by expression of *anxa4* (35) (**Fig. 1B**). Other top differentially expressed genes in cholangiocyte clusters included *lgals2b, tm4sf4, gramd1bb, gadd45bb*, and *klf6a* (**Fig. 1B, Table S1**). Cluster zChol2 uniquely showed a higher degree of overlap with hepatocyte gene expression patterns compared to both zChol1 and zAgr2+ (**Fig. 1B**). Interestingly, cluster zAgr2+ expressed the genes *agr2, icn*, and *tcnl*, which were found to be absent in the other cholangiocyte clusters (**Fig. 1B**). In human cholangiocytes, *AGR2* is expressed in both the tall epithelial cells covering large bile ducts as well as peribiliary gland cells, indicating that cluster zAgr2+ may identify a distinct population of cholangiocytes in zebrafish similar to these human *AGR2+* cells (36). Cholangiocyte populations were enriched for in GO processes including translation, apoptosis, and metabolism (**Fig. 1D**).

Endothelial cells and hepatic stellate cells (HSCs) grouped into a single cluster (zEC/HSC) which comprised approximately 7% of detected cells. Endothelial cells were identified by zebrafish markers *kdrl* (37) and *fli1a* (38) (**Fig. S1F-G**). Other enriched genes in this cluster included *socs3a, igfbp7, hspb1, fabp11a*, and *nppb* (**Table S1**). HSCs clustered with endothelial cells, identified by the expression of HSC marker *hand2* (39) and *colec11* (21, 22) (**Fig. 1B, S1H**).

Macrophages represented about 5% of all cells captured, identified by the expression of marker genes including *marco* and *grn1* (40, 41) (**Fig. 1B**). Two different clusters of macrophages were distinguished. First, an inflammatory macrophage cluster (zInf mac) was identified by the expression of inflammatory cytokines, including *il1b* and *cxcl19* (42) (**Fig. 1B**). These cells were enriched for known immune processes like respiratory burst and regulation of neutrophil migration by GO analysis (**Fig. 1E**). Second, a non-inflammatory macrophage cluster (zNon-inf mac) was found that notably lacked the expression of key inflammatory cytokines. GO analysis revealed that these cells were enriched for negative regulation of intrinsic apoptotic signaling in response to DNA damage and hemoglobin metabolic process (**Fig. 1E**).

Natural killer (NK) cells and T cells were also among the immune cell types captured by scRNA-seq of the adult zebrafish liver. These cells comprised about 3% of all analyzed cells and made up a single cluster (zNK/T). Key marker genes for these cell types included *il7r* (43, 44) and *trac* (44) (**Fig. 1B, S1I**). Other marker genes for cluster zNK/T included *sla2, dusp2, traf1, tnfrsf9b*, and *runx3* (**Fig. 1B, Table S1**). GO analysis on zNK/T cell markers revealed enrichment for immune pathways including regulation of neutrophil migration and regulation of intrinsic apoptotic signaling pathway (**Fig. 1E**).

Erythrocytes (zEry) represented roughly 3% of detected cells, and were identified by high expression of hemoglobin genes, including *ba1* and *hbaa1* (**Fig. S1J-K**) (45). Other top enriched genes in this cluster included *hbba1, hbba2, cahz, nt5c2l1*, and *epb41b* (**Fig. 1B, Table S1**). GO analysis revealed enrichment for carbon dioxide and gas transport in this cluster (**Fig. 1E**).

A cluster of neutrophils (zNeu) made up less than 1% of all cells in this dataset. Even in human single-cell transcriptomics, neutrophil subsets are difficult to capture due to low RNA content and the presence of endonucleases in the cell (46, 47). This zebrafish cluster was identified by the expression *mpx* (35) and *mmp13a* (40) (**Fig. 1B**). Other marker genes for the zNeu cluster included *si:dkeyp-75b4*.*10, mmp9*, and *itln1* (**Table S1)**. GO analysis revealed enrichment for processes including cellular response to antibiotics and regulation of leukocyte migration (**Fig. 1E**).

Finally, there was a small population of 111 cells (< 1% of all cells) that clustered separately from all other clusters (zApln+). This cluster was designated as zApln+ due to high expression of *apln*, a gene that was absent in all other captured cell types. Other marker genes for this cluster included *hbegfb, thbs1b, gp1bb, nfe2*, and *gcnt4a* (**Fig. 1B, Table S1**). Previous studies have shown that *apln* expression is highly expressed in endothelial tip cells, a type of endothelial cell found at the leading ends of sprouting vasculature that play a coordinating role in angiogenesis and vasculature organization in zebrafish (48, 49). Furthermore, *APLN* has been shown to be expressed in sprouting vasculature, particularly in the context of injury or disease, in mice (50), and humans (51, 52), indicating a conserved role for *APLN* across species.

This scRNA-seq dataset presented here provides an atlas for the distinct transcriptional profiles of the cell types that comprise the adult zebrafish liver and establishes an important resource for utilizing zebrafish as a tool to study liver function and disease.

### Key cell types and transcriptional profiles in the human liver are conserved in zebrafish liver

Since this is the first scRNA-seq dataset in adult zebrafish liver, we sought to determine the similarity of the adult zebrafish liver to the human liver single-cell transcriptome. We co-clustered our adult zebrafish liver single-cell dataset with a previously published human liver single-cell atlas (21). This human liver atlas contains 8,444 cells and is comprised of 20 distinct cell populations including hepatocytes, endothelial cells, cholangiocytes, and HSCs, as well as multiple immune cell clusters such as macrophages, T cells, NK cells, and B cells. Joint clustering of these datasets resulted in 21 clusters, again identified by calculating differential gene expression to create marker gene profiles for each cluster (**Fig. 2A-B, Table S3**).

**Figure 2.**
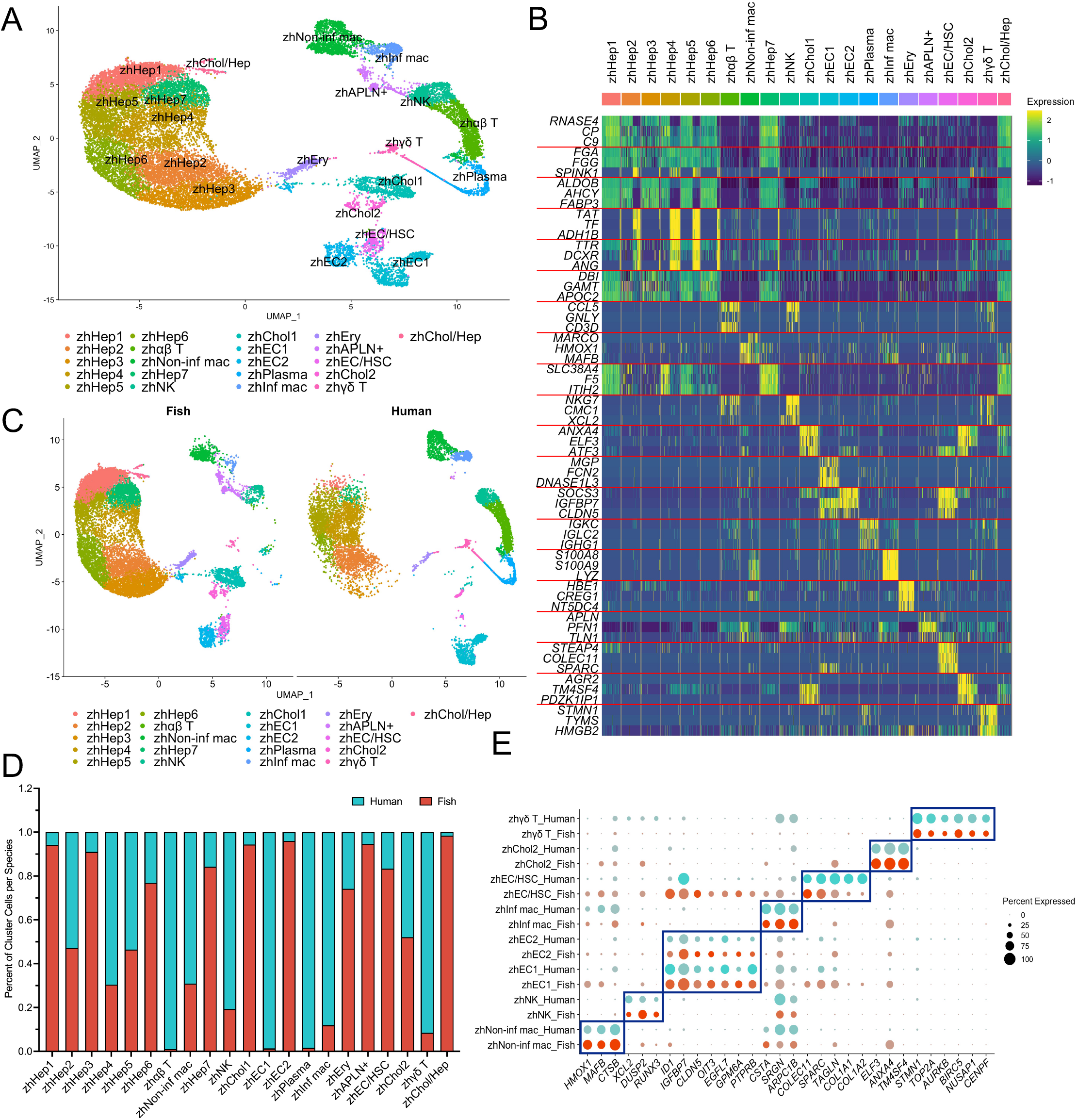
Joint clustering of adult human and zebrafish liver scRNA-seq. **A**. UMAP visualization of 21 clusters comprised of 8,444 adult human liver and 13,630 adult zebrafish liver cells. **B**. Heatmap of gene expression for three of the top differentially expressed genes in each cluster. **C**. UMAP of joint human and zebrafish clustering split by species. **D**. Bar graph showing the percentage of cells contributed to each cluster from each species. **E**. Dot plot showing differential expression of human cell type marker genes in select clusters split by species to show marker conservation. **Abbreviations**: zhHep – hepatocyte, zhαβ T – αβ T cell, zhNon-inf mac – non-inflammatory macrophage, zhNK – NK cell, zhChol – cholangiocyte, zhEC – endothelial cell, zhPlasma – plasma cell, zhInf mac – inflammatory macrophage, zhEry – erythrocyte, zhAPLN+ – *APLN+* cell, zhEC/HSC – endothelial cell/hepatic stellate cell, zhγδ T – γδ T cell, zhChol/Hep – cholangiocyte/hepatocyte mix.

Cells with like identities clustered together across species, as evidenced by the presence of both human and zebrafish cells in most clusters (**Fig. 2C-D, Table S4**). Hepatocytes accounted for the most clusters, generating 6 different joint populations (zhHep1-6). Human cholangiocytes largely clustered with the *agr2+* cholangiocyte (zhChol2) population from zebrafish, with very few human cholangiocytes found in the Chol1 cluster (**Fig. 2C-D**). Endothelial cell populations from human and zebrafish (zhEC1-2) largely clustered separately by species (**Fig. 2C-D**). Human and zebrafish HSCs clustered together in a single cluster, but zebrafish endothelial cell markers were also expressed in this cluster (zhEC/HSC) (**Fig. 2B-D**). Similar to analysis of the adult zebrafish scRNA-seq data alone, joint clustering also identified two macrophage populations, one inflammatory (zhInf mac) and one non-inflammatory (zhNon-inf mac) population, each composed of both human and zebrafish cells (**Fig. 2C-D**). Another interesting finding was that zebrafish NK/T cells showed the most co-clustering with human liver NK cells (zhNK), with equal proportions of the remaining cells clustering with either the αβ (zhαβ T) or γδ T cells (zhγδ T) (**Fig. 2C-D**). Finally, erythrocytes (zhEry) from both species also were found in a single cluster (**Fig. 2C-D**).

To further explore the similarity of human and zebrafish liver cell types involved in fibrosis, we compared expression of specific marker genes for human HSCs and other cell types that are known to interact with HSCs to regulate their activation (53). We found conservation of marker genes (21) for non-inflammatory macrophages (zhNon-inf mac) (*HMOX1, MAFB, CTSB*), NK cells (zhNK) (*XCL2, DUSP2, RUNX3*), endothelial cells (zhEC) (*ID1, IGFBP7, CLDN5, OIT3, EGFL7, GPM6A, PTPRB*), inflammatory macrophages (zhInf mac) (*CSTA, SRGN, ARPC1B*), HSCs (zhEC/HSC) (*COLEC11, SPARC, TAGLN, COL1A1, COL1A2)*, cholangiocytes (zhChol2) (*ELF3, ANXA4, TM4SF4)*, and γδ T cells (zhγδ T) (*STMN1, TOP2A, AURKB, BIRC5, NUSAP1, CENPF*) (**Fig. 2E**) across species, underscoring the similarities of cellular constitution between adult zebrafish and human livers.

Taken together, joint clustering of scRNA-seq data from adult human and adult zebrafish livers revealed a high degree of similarity in the transcriptional profiles and specificity of marker gene expression across species.

### Subclustering of zebrafish cluster zEC/HSC reveals distinct HSC and endothelial cell populations

HSCs are the key driver cell type in the development and progression of fibrosis (3), and we hypothesized that our scRNA-seq data could be used to better understand stellate cell biology *in vivo*. In our WT zebrafish liver atlas, endothelial cells and HSCs clustered together, and endothelial cells are known to have critical interactions with HSCs that regulate HSC activation (54). To identify other key cell types that interact with HSCs under physiological conditions, we utilized CellPhoneDB (55) to predict cell types that have receptor-ligand interactions with the zEC/HSC population from the WT zebrafish liver atlas (**Fig. 1A**). Immune cell clusters (zInf mac, zNon-inf mac, zNK/T, zNeu) and cluster zAgr2+ had the highest number of significant interactions with the zEC/HSC cluster (**Fig. S1L**), an important finding since immune cells and cholangiocytes are known to be important in regulating HSC activation in liver fibrosis in humans (53). Another cluster of interest is zApln+, which only interacted with the zEC/HSC cluster (**Fig. S1L**). With these interaction predictions, we focused on the immune, zAgr2+, and zApln+ populations as key zebrafish cell types that interact with HSCs.

To examine changes that occur in HSCs *in vivo* during liver fibrosis, we performed scRNA-seq on *mpi*^*+/mss7*^ adult zebrafish livers (N = 3). We previously reported that *mpi*^*+/mss7*^ adult zebrafish develop liver fibrosis, and that MPI-depleted HSCs have an activated phenotype (16), establishing the *mpi*^*+/mss7*^ adult zebrafish liver an excellent model to study mechanisms of HSC activation *in vivo*. To investigate the specific receptor-ligand interactions between zebrafish HSCs and other cell types, and to determine how these interactions are changed in HSCs in the context of liver fibrosis, we performed scRNA-seq on *mpi*^*+/mss7*^ adult zebrafish livers (N = 3). Sample viability ranged from 70 to 90% with samples contributing 4,169-5,742 cells (N = 15,286 cells). We clustered these samples with our WT zebrafish liver atlas in order to identify differences in cluster size and gene expression across genotype (**Fig. S2A, Table S5**). We then subset and reclustered the zEC/HSC cluster from the joint clustering of WT and *mpi*^*+/mss7*^ zebrafish liver samples to obtain a distinct HSC population we could interrogate. Using the predicted interactions of cell types with cluster zEC/HSC in our WT zebrafish liver atlas (**Fig. S1L**), we also subset and reclustered zInf mac, zNon-inf mac, zNK/T, zNeu, zAgr2+, and zApln+ clusters with cluster zEC/HSC from the WT and *mpi*^*+/mss7*^ zebrafish joint clustering to better investigate their reciprocal interactions with HSCs in physiological and fibrotic contexts. This strategy resulted in clusters of inflammatory macrophages (zInf mac), NK/T cells (zNK/T1 and zNK/T2), neutrophils (zNeu), non-inflammatory macrophages (zNon-inf mac), *agr2+* cholangiocytes (zAgr2+), and *apln+* cells (zApln+), as expected, but also allowed us to identify a distinct population of HSCs (zHSC), a mixed population of HSCs and endothelial cells (zEC-HSC), and 3 distinct endothelial cell populations (zEC1, zEC2, and zEC3) (**Fig. 3A**). All clusters contained both WT and *mpi*^*+/mss7*^ cells and were identified by marker genes determined by differential gene expression (**Fig. S2B-C**). The zHSC cluster represented roughly 0.1% of all cells captured in the total liver dataset and was identified by *hand2* expression (39), which was specifically expressed in the zHSC and zEC-HSC clusters but only captured in a small percentage of cells (**Fig. S2D**). Top differentially expressed genes for this cluster included *col1a1b, ifitm1, ccl25b, sparc, steap4*, and *colec11* (**Fig. 3B, Table S6**). *mpi*^*+/mss7*^ livers did have a higher percentage of captured HSCs on average, although this difference was not significant (**Fig. 3C, Table S7**).

**Figure 3.**
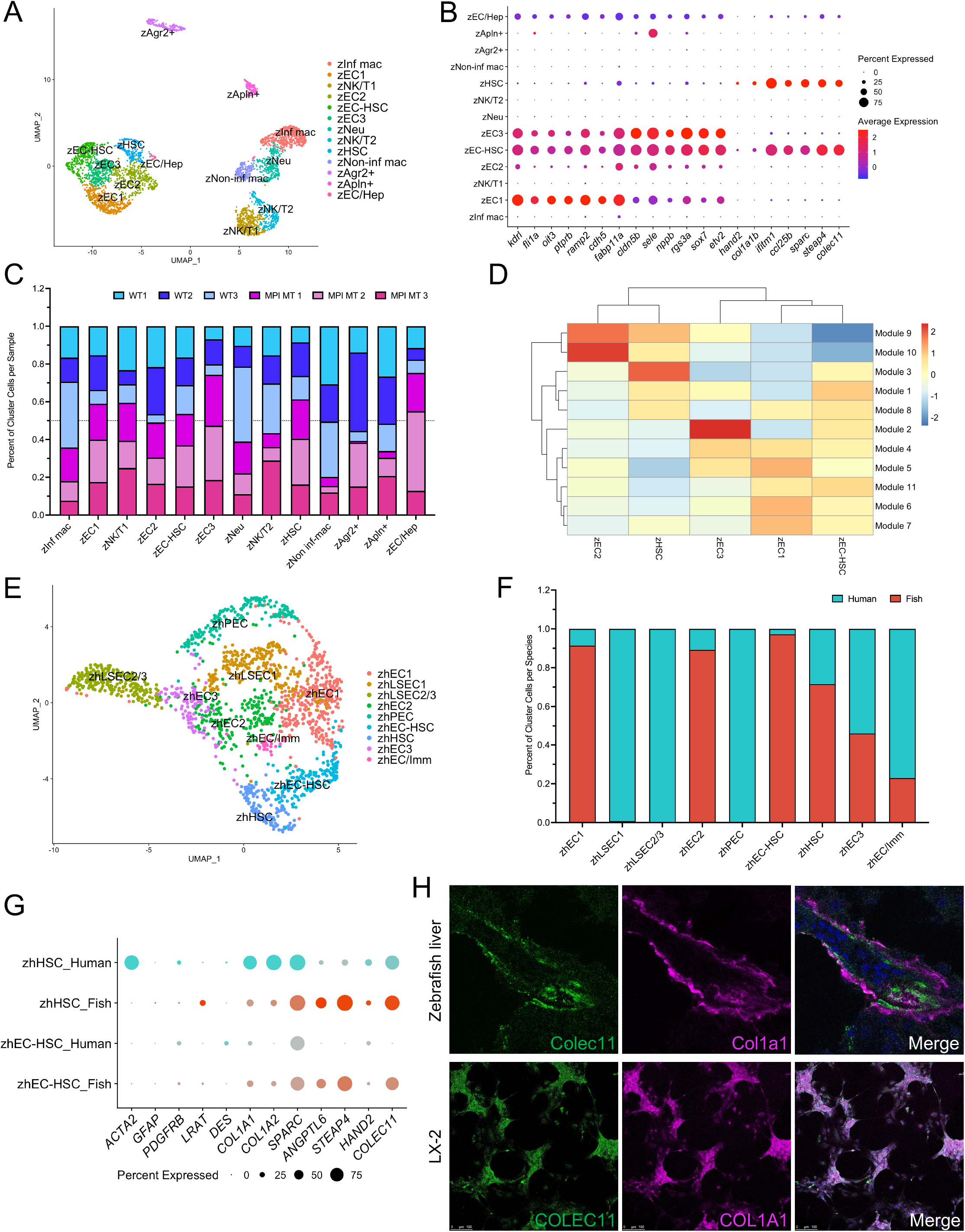
Dissecting out zebrafish HSCs and endothelial cells. **A**. UMAP visualization of 13 clusters comprised of WT and *mpi* ^*+/mss7*^ adult zebrafish liver cells subset from total liver cell clustering. **B**. Dot plot of gene expression for top differentially expressed genes in clusters zEC1, zEC2, zEC3, zEC-HSC, and zHSC. **C**. Bar graph showing the percentage of cells contributed to each cluster from each sample. **D**. Heatmap showing expression of modules of co-regulated genes correlating with endothelial cell and HSC clusters. **E**. UMAP visualization of 9 clusters comprised of adult human liver and adult zebrafish liver endothelial cells and HSCs. **F**. Bar graph showing the percentage of cells contributed to each cluster from each species. **G**. Dot plot showing differential expression of HSC marker genes split by species to show marker conservation. **H**. Immunofluorescent staining for COLEC11 and COL1A1 on WT adult zebrafish liver cryosections (imaged at 63X) and LX-2 cells (imaged at 20X). **Abbreviations**: **zebrafish clustering:** zInf mac – inflammatory macrophage, zEC – endothelial cell, zNK/T – NK cell/ T cell, zEC-HSC – endothelial cell-hepatic stellate cell doublet, zNeu – neutrophil, zHSC – hepatic stellate cell, zNon-inf mac – non-inflammatory macrophage, zAgr2+ – *agr2+* cell, zApln+ – *apln+* cell, zEC/Hep – endothelial cell/hepatocyte mix; **zebrafish and human joinit clustering:** zhEC – endothelial cell, zhLSEC – liver sinusoidal endothelial cell, zhPEC – portal endothelial cell, zhEC-HSC – endothelial cell-hepatic stellate cell doublet, zhHSC – hepatic stellate cell, zhEC/Imm – endothelial cell/immune cell mix.

Three distinct endothelial cell clusters (zEC1, zEC2, and zEC3) were also identified, collectively making up approximately 4% of all captured cells from total liver clustering and identified by expression of the known endothelial cell marker, *kdrl* (37) (**Fig. S2E**). Top differentially expressed genes for endothelial cell clusters included *oit3, ptprb*, and *ramp2* for zEC1, and *cldn5b, sele*, and *nppb* for zEC2 and zEC3 (**Fig. 3B, Table S6**).

Liver sinusoidal endothelial cells (LSECs) are known to physically interact with HSCs (56). In our dataset, we identified a cluster present in all WT and *mpi*^*+/mss7*^ samples that demonstrated expression for genes that were enriched in both endothelial cell and hepatic stellate cell clusters (zEC-HSC) (**Fig. 3B**). Furthermore, the cells in this cluster had a higher number of detected UMIs on average than in the zHSC or zEC clusters, indicating that this population may be made up of endothelial cell and HSC doublets (**Fig. S2F**).

The human liver has unique endothelial cell types (21) and LSECs are known to play a critical role in the regulation of HSC activation (54). Thus, we sought to further characterize the different zebrafish endothelial cell clusters in our dataset to identify specific zebrafish endothelial subtypes involved in HSC regulation. To further validate that clusters zEC1, zEC2, and zEC3 are representative of unique populations of zebrafish endothelial cells with distinct transcriptional profiles, we used an additional and alternative method for determining differential gene expression across the endothelial cells and HSCs. The pseudotime analysis tool, Monocle, employs a method for analyzing differential gene expression of cells in UMAP space using a statistic from Moran’s *I*, a type of spatial autocorrelation analysis (57–62). Rather than identifying differentially expressed genes between the defined clusters, this analysis identifies genes that are expressed in specific focal regions of cells in UMAP, which can then be used to create modules of gene co-expression that are agnostic of cluster identity. The expression of each of these modules can then be visualized in each of the clusters in the analyzed dataset. We subset the zebrafish WT endothelial cell and HSC populations (**Fig. S3A**) and performed this analysis to identify differentially expressed genes across these cells in UMAP space (**Table S8**), categorized into modules of co-expressed genes (**Table S9**). This revealed unique expression of modules for all 3 clusters of endothelial cells, as well as the zHSC cluster, indicating that these populations are transcriptionally distinct from one another (**Fig. 3D**). Furthermore, numerous modules specifically-expressed by the zHSC and endothelial cell clusters were also expressed by the zEC-HSC cluster. The shared expression of these modules indicates transcriptional similarity of the zEC-HSC cluster to both zHSC and endothelial cell clusters, further implying that these cells are likely doublets of HSCs and endothelial cells (**Fig. 3D**). Distinct modules were then confirmed to have cluster-specific expression by validating increased expression of the genes comprising each module across the 5 clusters (**Fig. S3B-C**). These results confirm that the zebrafish endothelial cell clusters in our dataset have significantly different transcriptional profiles, indicating that the zebrafish liver contains distinct subtypes of endothelial cell populations with unique functions similar to human liver (21).

### Demonstrating conservation of HSC gene expression with human and zebrafish joint clustering

The use of adult zebrafish to study liver fibrosis has been limited thus far. Therefore, we sought to determine the conservation of endothelial and stellate cell populations between zebrafish and human livers. WT zebrafish endothelial cell and HSC populations were jointly clustered with human HSCs, LSECs, and portal endothelial cells. This resulted in 3 distinct human endothelial cell populations (zone 1 LSECs (zhLSEC1), zone 2/3 LSECs (zhLSEC2/3), and portal endothelial cells (zhPEC)), recapitulating the populations identified in MacParland et. al. (21) (**Fig. 3E-F, Table S10-11**). There were 3 other endothelial cell populations, two of which were comprised primarily by zebrafish cells (zhEC1 and zhEC2) and one made of equal proportions of human and zebrafish cells (zhEC3) (**Fig. 3E-F, Table S10-11**). Overall, zebrafish and human endothelial cells did not cluster together due to poor conservation of human endothelial cell marker gene expression in zebrafish (**Fig. S4A-B**).

In contrast to the ECs, the majority of human and zebrafish HSCs clustered into a single cluster (zhHSC), with the exception of the zebrafish EC-HSC doublets (zhEC-HSC) (**Fig. 3E-F, S4A**). Expression of human HSC marker genes *ACTA2, COL1A1, COL1A2, SPARC*, and *COLEC11* (21, 63) was observed in over 75% of all human HSCs (**Fig. 3G**). Of these genes, *SPARC* and *COLEC11* were highly captured in zebrafish HSCs, while zebrafish marker *HAND2* and markers determined from our zebrafish scRNA-seq dataset, *ANGPTL6* and *STEAP4*, were poorly captured in human HSCs (**Fig. 3G**). Commonly used protein markers for human HSCs, including GFAP, PDGFRB, LRAT, DES (63, 64), were poorly captured at the transcriptional level by scRNA-seq in both human and zebrafish HSCs (**Fig. 3G**). These findings indicate that, at the transcriptional level, traditional markers of human and zebrafish HSCs show lower levels of conservation. We utilized our scRNA-seq data to identify additional HSC markers in zebrafish. We found Colec11 to be highly expressed in human and zebrafish HSCs (**Fig. 3G**) and tested its expression and specificity for zebrafish HSCs by immunofluorescence (IF) staining on liver tissue sections from WT adult zebrafish. We were able to identify Colec11-expressing cells and demonstrated colocalization with Col1a1 (**Fig. 3H**). We also performed IF staining for COLEC11 and COL1A1 on human hepatic stellate cells (LX-2s) and found that COLEC11 and COL1A1 were co-expressed by these cells (**Fig. 3H**). To our knowledge, this is the first use of Colec11 as a novel marker for zebrafish HSCs and demonstrates that COLEC11 expression is conserved across human and zebrafish HSCs.

Taken together, while human and zebrafish endothelial cells have distinct gene expression patterns, human and zebrafish HSCs show conservation in transcriptional profiles and marker gene expression, with Colec11 demonstrating specific expression in HSCs in zebrafish. The similarity of human and zebrafish HSCs strengthens the case for using the zebrafish to study HSC activation and liver fibrosis.

### Key pathways in HSC activation, angiogenesis, and immune response are altered in MPI-depleted HSCs

To identify key canonical pathways and upstream regulators that are altered in the context of fibrosis, we used Ingenuity Pathway Analysis (IPA) on the differentially expressed genes (DEGs) between *mpi*^*+/mss7*^ and WT HSCs from our zebrafish adult liver scRNA-seq datasets (**Fig. 3A, Table S13**). The hepatic fibrosis signaling pathway was significantly activated in *mpi*^*+/mss7*^ HSCs further supporting that loss of Mpi activates HSCs and promotes liver fibrosis. Other canonical pathways known to be involved in liver fibrosis and HSC activation, including IGF-1, estrogen receptor, and mTOR signaling, were also found to be significantly altered (65–67). Furthermore, TNF, TGFβ-1, IL-1β, and EGF were among the most significantly activated upstream regulators and have all been demonstrated to be key players in the activation of HSCs and the progression of liver fibrosis (68–71). These data from *in vivo* zebrafish *mpi*^*+/mss7*^ HSCs are consistent with an activated phenotype and likely contribute to the dense fibrosis observed *in vivo* (16).

To determine whether these predicted pathway alterations found in zebrafish HSCs are similar to the effects of MPI-depletion in human HSCs, we performed IPA on DEGs collected from bulk RNA-seq data from two models of human MPI-depleted HSCs: a stable *MPI* mutant LX-2 line (**Table S14**), and TWNT-4 cells with acute depletion of MPI by siRNA (siMPI) (**Table S13**). We created the stable *MPI* mutant LX-2 line using CRISPR/Cas9 with a gRNA targeting exon 3. This created a 39 base pair deletion in exon 3 (NC_000015.10:g.74891387_74891425) (**Fig S5A**) and resulted in a deletion-insertion mutation replacing MPI amino acids 51-64 with an isoleucine (NP_001276084.1:p.(Met51_Asn64delinsIle) (**Fig. S5B**). *MPI MT* LX-2 cells had a residual MPI activity of 21.4% compared to WT controls **(Fig. 4A**), and decreased amounts of MPI protein (**Fig. 4B**). *MPI MT* LX-2 cells also demonstrated elevated COL1A levels indicative of an activated phenotype (**Fig. 4B**). We also analyzed HSCs with acute depletion of MPI; siMPI-treated TWNT-4s had a residual MPI activity of 44.4% compared to controls (**Fig. S5C**). We compared significance scores for all altered pathways in IPA, and found that conserved significantly-altered pathways in MPI-depleted HSCs among zebrafish and humans were similar across datasets. These included pathways involved in HSC activation, angiogenesis, and immune response, all of which are known to play key roles in the pathogenesis of liver fibrosis (**Fig. 4C**) (53). We also compared the activation z-score of upstream regulators predicted to be altered across all three datasets and found many significant predicted alterations in the activity of upstream regulators, including EGF, FGF, ERK1/2, and Pdgf (complex), which are known to be key signaling pathways in HSC activation (72–75) (**Fig. 4D**). Taken together, these data indicate that MPI-depletion in zebrafish and human HSCs, *in vivo* and *in vitro*, lead to similar and significant alterations in fibrogenic pathways, further demonstrating the power of zebrafish HSCs to recapitulate human HSC phenotypes.

**Figure 4.**
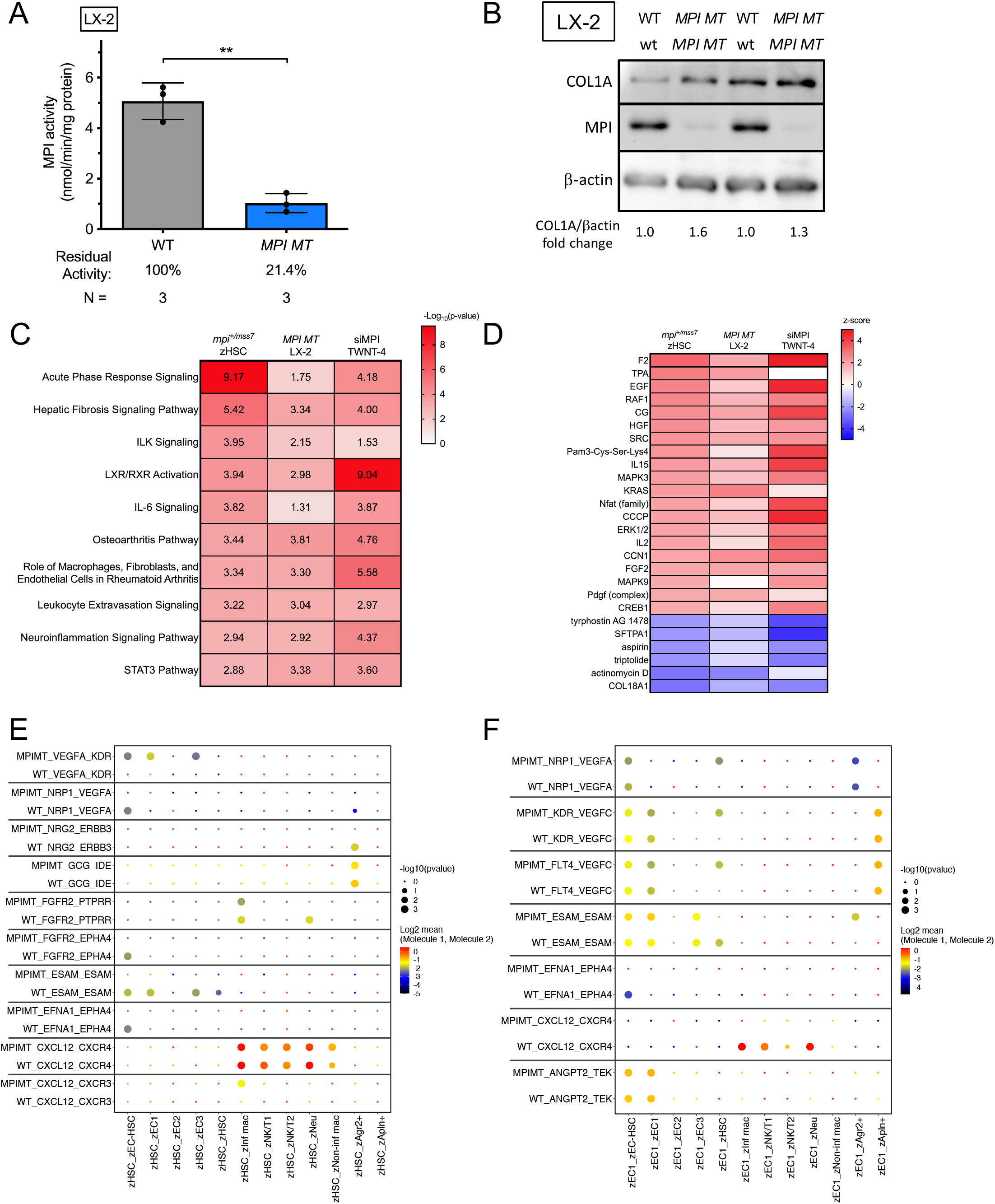
Transcriptional profiling and cell-cell interactions of MPI-depleted HSCs. **A**. Bar graph showing MPI enzymatic activity in WT and *MPI MT* LX-2 cells. **B**. Western blot for MPI and COL1A1 WT and *MPI MT* LX-2 cells. **C-D**. Ingenuity Pathway Analysis conducted on differential gene expression from 3 datasets: *mpi* ^*+/mss7*^ zebrafish single cell HSCs (scHSCs), MPI mutant LX-2 HSCs, and siMPI TWNT-4 HSCs. Heatmaps display (**C**) p-values for canonical pathway activity alteration and (**D**) z-score of activation/inhibition of upstream regulators. **E-F**. Dot plot of receptor-ligand interaction scores determined by CellPhoneDB analysis of scRNA-seq gene expression in WT and mpi ^*+/mss7*^ zebrafish (**E**) zHSC and (**F**) zEC1.

We next examined specific receptor-ligand pairs that are expressed by zebrafish HSCs and their interacting partners *in vivo* to identify altered interactions that may play a role in HSC activation and fibrosis. We conducted CellPhoneDB (55) analysis on the HSC subset reclustering of our zebrafish liver scRNA-seq data (HSCs, endothelial cells, immune cells, *agr2*+ cells, and *apln*+ cells) (**Fig. 3A**) for both WT and *mpi*^*+/mss7*^ mutants. CellPhoneDB predicted significant receptor-ligand interactions to exist between HSCs and endothelial cells, macrophages, NK/T cells, neutrophils, and *agr2*+ cholangiocytes. Interestingly, in the *mpi*^*+/mss7*^ mutants, cluster zHSC was predicted to have a significant interaction via VEGFA-KDR with both zEC1 and zEC3, which was absent in the WT dataset (**Fig. 4E**). VEGF is a critical ligand for the maintenance of the vasculature and has been show to play a key role in the activation of HSCs (53, 76, 77). Furthermore, multiple interactions between the WT zHSC and zEC-HSC cluster cells were lost in the *mpi*^*+/mss7*^mutants (**Fig. 4E**). We also investigated differences in the expression of receptor-ligand pairs in the endothelial cell populations and found that significant interactions were predicted for NRP1-VEGFA, KDR-VEGFC, and FLT4-VEGFC in zEC1 interaction with zHSC in the *mpi*^*+/mss7*^mutants but not in the WTs (**Fig. 4F**). These data highlight differences in receptor-ligand interactions between WT and *mpi*^*+/mss7*^ cells, indicating that key interactions between HSCs, endothelial cells, and immune cell types are disrupted with loss of Mpi, which may play a role in the activation of HSCs.

## Discussion

The public health burden of cirrhosis continues unchecked with an unmet need for effective anti-fibrotic therapeutics. Furthermore, the mechanisms regulating liver fibrogenesis are extremely complex and include orchestration of multiple cell-types and cell-cell interactions (78, 79). Here, we demonstrate the potential for expanding the use of scRNA-seq in the adult zebrafish as a tool towards identifying key cell-specific roles in liver fibrosis which may be candidates for therapeutic intervention.

We have created the first single-cell atlas of the adult zebrafish liver, comprised of transcriptionally unique populations of hepatic cell types, including hepatocytes, biliary cells, endothelial cells, immune cells, and HSCs. When choosing an animal model to study a human disease, the degree of conservation for specific cell-types is critical. Although conventional studies have shown that the zebrafish liver is similar to the human liver (10), our comparative analysis using scRNA-seq reveals highly conserved marker genes in like cell types with similar transcriptional profiles indicative of shared identity and functional roles in the liver. While the zebrafish liver is structurally different from the human liver and not known to have architectural zonation (10, 33, 34), our zebrafish liver scRNA-seq dataset revealed three main, but distinct, groups of hepatocytes enriched for pathways involved in oxidative phosphorylation (zHep1, zHep7, and zHep8), cholesterol, steroid, and lipid metabolism (zHep2, zHep3, zHep4, and zHep5), and fatty acid transport and glucose metabolism (zHep6). Whether these differences are related to the spatial location of these hepatocytes in the zebrafish liver or indicate zonation of the zebrafish liver similar to that of the human liver will require further study and emerging spatial scRNA-seq platforms may be applicable (80).

Given that HSCs are the primary cell type in liver fibrosis (2), conservation of this cell type in zebrafish is critical for establishing the adult zebrafish as an effective model for studying liver fibrosis. We found that zebrafish HSCs demonstrate a high similarity to human HSCs. This is seen through both co-clustering and conservation of marker genes across species. Furthermore, there are limited tools that currently exist to identify and study HSCs in the zebrafish liver. Only *Tg(hand2:GFP)* (39) zebrafish and GFAP IF staining (81) have been shown to properly identify HSCs in zebrafish. Our work expands upon this by identifying a new marker for zebrafish HSCs that could be used to both visualize HSCs *in vivo* and isolate HSCs from the zebrafish liver. To our knowledge, this is the first study to demonstrate that Colec11 is one such marker that shows positive staining of zebrafish HSCs.

Endothelial cells have established roles in both activation and reversion of HSCs in fibrogenesis (54). The human liver is known to have different endothelial cell types, including LSECs and portal endothelial cells (21), but little is known about whether zebrafish liver endothelial cells are a homogenous or heterogenous population. Our data revealed multiple endothelial cell populations characterized by unique modules of gene expression, demonstrating that discrete populations of endothelial cells do exist within the zebrafish liver. However, conclusions are limited as human marker genes that discriminate between different endothelial cell populations do not show the same specificity for zebrafish endothelial cells. Further studies are needed to explore how these zebrafish endothelial cell subpopulations differ from one another in spatial location and function.

We have previously shown that loss of MPI induces HSC activation and liver fibrosis (16). Pathway analysis comparing three transcriptomic datasets across zebrafish and human HSCs, one from *in vivo mpi*^*+/mss7*^ zebrafish HSCs and two from *in vitro* MPI-depleted human HSC lines, showed alteration in similar pathways, demonstrating conserved response to loss of MPI. Significant alteration of the hepatic fibrosis signaling pathway highlights the role of MPI in the regulation of HSC activation. Other altered pathways implicate specific endothelial cell and immune cell processes as being involved in HSC activation and fibrosis. Furthermore, we identified receptor-ligand interactions between HSCs and interacting partner cell types in the zebrafish liver, as well as notable differences in these receptor-ligand pairs with Mpi depletion. Interestingly, significant interaction between VEGF ligands and their receptors was predicted between HSCs and endothelial cells in *mpi*^*+/mss7*^ cells but not WT cells, supporting a link between MPI loss and VEGF signaling, and consistent with angiogenic defects found in MPI knockout mice (82). VEGF signaling is a key regulator of endothelial cells and plays a role in liver fibrosis (53, 76, 77), thus using our model to explore these interactions will provide greater understanding of how these signaling pathways are involved in fibrosis, and also identify the role they play in liver fibrosis in the context of MPI depletion. These studies can lay the foundation for future modeling of human fibrotic liver diseases using zebrafish.

In summary, the use of scRNA-seq in zebrafish livers highlights the similarities and differences to human liver, supports its use as a valuable tool to study liver fibrosis, identifies new tools to study HSCs, and lends insights into important *in vivo* interactions for further investigation.

## Materials and Methods

### Zebrafish maintenance

Procedures were performed in accord with the Icahn School of Medicine at Mount Sinai (ISMMS) Institutional Animal Care and Use Committee. Adult fish were maintained on a 14:10 light/dark cycle at 28°C. AB WT and the *mpi* ^*mss7*^ (15) zebrafish strain were used.

### scRNA-seq on adult zebrafish livers

This protocol was adapted from Nayar et al., 2021 (24).

### Single-cell suspension

Whole livers were dissected from 18-month-old adult, male zebrafish that were anesthetized in Tricaine (1.25g per 500 ml water, pH 7.4) and then euthanized in ice-cold water for 15 minutes. Samples were kept at RT throughout preparation unless otherwise stated. Livers were collected in 2 mL autoMACS Rinsing Solution + 0.5% BSA (Miltenyi Biotec). Tubes were then rocked for 30 min at room temperature to reduce epithelial cell proportions, followed by centrifugation at 4,200 rpm for 5 min. After removal of the supernatant, liver pellets were resuspended in 2 mL PBS + Ca^2+^ + Mg^2+^ + 1 mg DNaseI + 1 mg collagenase IV and rocked for 35 min at 37 °C at 100 rpm. Mixtures were filtered through a 70-μm filter into a new tube and spun at 4,200 rpm for 5 min. Cell pellets were resuspended in a final volume of 300 μL PBS + 0.5% BSA. Cells were counted with AO/PI dye using the Nexcelom Biosciences cell counter. Ten thousand cells were loaded onto the 10X Genomics Chromium Controller instrument within 30 min of completion of cell suspension using GemCode Gel Bead and Chip (10X Genomics). Cells were partitioned into gel beads in emulsion in the controller in which cell lysis and reverse transcription occurred. Libraries were prepared using 10X Genomics library kits and sequenced on Illumina NextSeq500, according to manufacturer’s recommendations.

### Alignment, transcriptome assembly and quality control

Raw BCL (raw base call) files generated from Illumina were demultiplexed using cellranger v.4.0.0, which uses the bcl2fastq pipeline, into single-cell FASTQ files. FASTQ files were then aligned to the Ensembl GRCz11 zebrafish genome and transcript count matrices were generated using default parameters in cellranger count. The raw (unfiltered) output matrices were then used for the clustering and downstream analysis in the R package Seurat v4.0 (25– 28). Density plots for unique molecular identifiers (UMIs), number of genes, and mitochondrial percentages were analyzed for each sample. Data were filtered to include cells with > 200 UMIs, 200 > genes per cell > 3000, and < 50% mitochondrial transcripts. Samples were individually normalized, and the 2000 most variable genes were identified for each sample. ‘FindIntegrationAnchors’ was used to integrate all zebrafish samples. The data were scaled in Seurat, and dimensionality reduction was then performed; the first 15 principal components were used to generate clustering, with a resolution of 0.8. Enriched genes for each cluster were identified using the ‘FindAllMarkers’ function. The default Wilcoxon Rank Sum test was used to determine significance, and we employed a cutoff of Log_2_(fold change) = 0.25.

### Differential gene expression on scRNA-seq data

To determine differential gene expression between clusters and samples, the ‘FindMarkers’ function in Seurat v4.0 was used (25–28). The default Wilcoxon Rank Sum test was used to determine significance.

### Gene ontology (GO) enrichment analysis

This protocol was adapted from Willms et al., 2020 (83). Marker genes (p-value < 0.05) for each cluster were analyzed in GOrilla (Gene Ontology enrichment anaLysis and visuaLizAtion tool) to determine significantly enriched GO processes (84). Genes were analyzed in a two-list unranked comparison using the list of all genes present in the dataset as the background set. REVIGO (Reduce and Visualize Gene Ontology) was used to measure semantic similarity using the SimRel analysis and a cutoff value of 0.4 for allowed similarity (85). The three most significantly enriched processes are shown for each cluster. Plots were created using Prism 9.

### Human and zebrafish liver integrated clustering

A list of human-to-fish orthologs was generated using ZFIN zebrafish–human orthologs database (20 August 2020) (24). The orthologs in the database were curated considering three factors: conserved genome location, amino acid sequence comparison and the phylogenetics tree. Zebrafish gene names were converted to the orthologous human gene name using the ZFIN ortholog database. Filtered human liver scRNA-seq data from five healthy, neurologically deceased donors was obtained from https://github.com/BaderLab/HumanLiver (21). Joint clustering of human and zebrafish data was performed using Seurat v4.0 (25–28). The 2000 most variable genes were determined, and the datasets were integrated using ‘SelectIntegrationFeatures’ and ‘FindIntegrationAnchors’. The first 15 principal components were used to perform clustering at a resolution of 0.8.

For joint clustering of HSCs and endothelial cells only, these cell types were first subset from the human and zebrafish (gene names converted to human orthologs) datasets respectively, and then the same workflow was applied, using the first seven principal components and a resolution of 0.8, to generate clustering.

### Identifying ligand-receptor interactions

Zebrafish gene names were converted to the orthologous human gene name as described in ‘Human and zebrafish liver integrated clustering’. CellPhoneDB v.2.0 (55) was run using its statistical analysis method with normalized counts data and cell annotation input files from each zebrafish scRNA-seq dataset. For **Fig S1G**, the WT zebrafish liver scRNA-seq transcriptome atlas was used for analysis. For **Fig 4E-F**, WT and *mpi* ^*+/mss7*^, cells were split by genotype into individual datasets and then analyzed using CellPhoneDB v.2.0. Heatmaps displaying the log-count of interactions were created using ‘heatmap_plot’. Dot plots of specific ligand-receptor interactions between cell type pairs were created using the R script for the ‘dot_plot’ function and modified ligand-receptor p-value and mean spreadsheets that contained data for both genotypes. The ‘dot_plot’ R script was obtained from https://github.com/Teichlab/cellphonedb.

### HSC and HSC-interacting cell types subclustering

WT and *mpi* ^*+/mss7*^ samples were clustered together using the same workflow described in ‘scRNA-seq on adult zebrafish livers’. Once the data were clustered and cell types were annotated, HSCs other cell types of interest were subset from the total dataset. The data were then normalized, and variable features were identified. After dimensionality reduction, the first 15 principal components and a resolution of 0.8 were used to generate clustering.

### Identifying modules of co-regulated genes in endothelial cell populations

The ‘graph_test’ function in Monocle 3 (57–62) was used to identify genes that are variable in expression across zebrafish WT endothelial cell and HSC clusters. ‘find_gene_modules’ was then used to perform UMAP analysis on significant genes (q-value ≤ 0.05) and group the genes into modules by using Louvain community analysis. The ‘aggregate_gene_expression’ function was then used to aggregate the expression of all genes in each module to visualize the expression of each module in each cluster.

### Immunofluorescence staining

Adult zebrafish livers were fixed in 4% paraformaldehyde (PFA) in phosphate-buffered saline (PBS) overnight immediately following dissection. After fixation, livers were incubated in increasing concentrations of sucrose in PBS (10%, 20%, and 30%) for 24 hrs each. Livers were embedded in optimum cutting temperature (OCT) compound (Tissue-Tek), and 10μm serial sections were obtained using the Leica CM3050 S Research Cryostat. Sections were washed in PBS + 0.1% Tween-20 (PBST). Tissue sections were blocked with 5% fetal bovine serum (FBS) and 2% bovine serum albumin (BSA) in PBST for 2 hours at room temperature (RT). LX-2 cells were seeded at 200,000 cells/mL on chamber slides (Nunc Lab-Tek; Thermo Fisher), grown overnight, and fixed in 4% PFA for 15 minutes. Fixed cells were permeabilized with PBS + 0.4% Triton X 100, and blocked with PBS + 0.25% Triton X 100 + 5% FBS + 2% BSA for 1.5 hours at RT. Sections/cells were stained with 1:100 COLEC11 antibody (ab278063; Abcam) and 1:100 COL1A1 antibody (zebrafish sections: ab23730, Abcam; human cells: bs-10423R; Bioss) overnight at 4°C and then with 1:250 goat anti-mouse Alexa Fluor 488 (A11008; Life Technologies, Waltham, MA) and 1:250 goat anti-rabbit Alexa Fluor 647 (A21244; Life Technologies, Waltham, MA) for 1.5 hours in the dark at RT. Sections were mounted with ProLong Diamond Antifade Mountant with 4′,6-diamidino-2-phenylindole (DAPI; Life Technologies) and imaged using a Leica SP5 DMI (zebrafish sections: 63×; human cells: 20×). Imaging was conducted in the Microscopy CoRE at the Icahn School of Medicine at Mount Sinai.

### Cell culture

LX-2 (86) and TWNT-4 (87) human HSC lines were maintained in complete medium (Dulbecco’s modified Eagle’s medium [DMEM]) supplemented with 10% FBS and L-glutamine, and penicillin/streptomycin) and routinely tested for mycoplasma using the Venor GeM Mycoplasma Detection Kit (Sigma-Aldrich). For MPI knockdown experiments, small interfering RNAs (siRNAs) targeting human MPI were transfected using Lipofectamine RNAiMAX transfection reagent (ThermoFisher, Waltham, MA), as previously described (15). Cells were collected 48 hours after transfection for total RNA or protein.

### CRISPR-Cas9 mutagenesis of *MPI* in human HSCs

To generate a stable mutant MPI HSC line, the LX-2 human HSC line was infected with lentivirus harboring plasmid sequences from the pLV-U6g-EPCG expression vector (CRISPRV, MilliporeSigma). Approximately 2,000 cells were infected at MOI 50 in DMEM + 10% FBS and 4μg/ml polybrene. The lentivirus harbored both gRNA targeting the start codon of MPI (5’-ATGGGGACTCACCCCCGAG-3’), and a Cas9 open reading frame flanked by GFP and puromycin-resistance elements. Transduced cells with positive integration were selected for 3 days with puromycin (2μg/ml), expanded, and confirmed by sequencing (Primers: F: 5’-AACTCAGGGTGGCAGGTTTC −3’; R: 5’-AAAGTCACGGGAGGGCCTTA-3’) (**Fig. S5B**).

### MPI enzyme activity assay

MPI activity assay was performed on mammalian cell protein extract according to our published protocol (15, 88). Briefly, cell lysates were homogenized and MPI activity was assessed in 15 ug of protein extract using a coupled-fluorometric assay.

### Bulk RNA-seq on human HSC cell lines

LX-2 and TWNT-4 human HSCs were lysed in TRIzol (Life Technologies) and total RNA was purified following the supplied protocol. For LX-2 control and *MPI MT* preparations, samples were subsequently treated with DNAse and further purified using the Quick RNA microprep kit (Zymo research, R1050) following the supplied protocol. RNA quality check, library preparation and sequencing (RNA-seq) was performed by Genewiz (South Plainfield, NJ). Briefly, mRNA was enriched by poly(A) selection, fragmented cDNA synthesized by random priming. 5’-phosphorylated and dA-tailed library was ligated to adapter, PCR enriched and sequenced using Illumina HiSeq 2×150. Reads were trimmed using Trimmomatic v.0.36, and trimmed reads were mapped to the Homo sapiens GRCh38 reference genome available on ENSEMBL using the STAR aligner v2.5.2b. Unique gene hit counts were calculated using featureCounts from the Subread package v.1.5.2, and differential gene expression analysis was performed using DESeq2 by Genewiz (South Plainfield, NJ) (89, 90).

For TWNT-4 HSC transfected with non-targeting control dsiRNA (NC) or dsiRNA targeting MPI (siMPI) samples, RNA quality was determined with the Agilent 2100 BioAnalyzer. cDNA libraries were prepared using the Illumina TruSeq RNA sample preparation version 2 protocol, DNA quality was checked with the Agilent 2100 BioAnalyzer. and libraries were sequenced with PE 100bp on the Illumina platform HiSeq2500. Trimmed reads by Trimmomatic (DOI: 10.1093/bioinformatics/btu170) were mapped to the Ensembl human reference genome GRCh38 using HISAT2 with default parameters. To estimate gene expression, reads mapped at exons of ensemble annotated genes were calculated with the ‘union’ and ‘no stranded’ mode with by HTSeq (https://doi.org/10.1093/database/baw093, doi:10.1093/bioinformatics/btu638). DESeq2 in Bioconductor (89, 90) was adopted to test differential gene expression. Adjusted p-value (FDR) < 0.05 was treated as significant.

### Ingenuity Pathway Analysis (IPA) on differential gene expression

Differential gene expression was calculated for *mpi* ^*+/mss7*^ vs. WT HSCs as described in ‘Differential gene expression on scRNA-seq data’. Differentially expressed genes, log fold changes, and p-values were imported into IPA for all three datasets. Data were analyzed through the use of IPA (QIAGEN Inc., https://www.qiagenbioinformatics.com/products/ingenuitypathway-analysis (91). Core analysis was conducted on each dataset using log fold change values. Figures were generated using Prism 9.

## Code availability

All R codes used for data analysis and visualization have been uploaded to https://github.com/jkmorrison/zebrafish-liver. The ZFIN genomic databases can be found at https://zfin.atlassian.net/wiki/spaces/general/pages/1891412257/Genomic+Resources+for+Zebrafish.

## Acknowledgements and funding sources

We thank the Human Immune Monitoring Center at ISMMS, including L. Walker and H. Stefanos, for running scRNA-seq on our zebrafish samples. We thank the Microscopy CoRE for their technical assistance. We thank Nataly Shtraizent, Kirsten Sadler Edepli, and the NYU Abu Dhabi Sequencing Core for their assistance with bulk RNA-seq on human HSCs.

This research was funded by R01 DK121154, R01 DK121154-01A1S1, and the AASLD Foundation Bridge Award (to J.Chu) and R01 DK123758 (to J.H.Cho).

## Author contributions

J.K.M. and J.C. conceived the project. J.M., C.D., I.L.A., S.N., M.G., J.H.C. and J.C. designed the experiments, and J.M., C.D., I.L.A, and C.Z. performed them. J.M. and J.C. wrote the manuscript, with discussions and edits from J.M., C.D., I.A., S.N., J.H.C.. J.M., C.D., I.L.A., S.N., J.H.C., and J.C. provided intellectual input throughout the study’s progression. M.G. and C.Z. provided assistance with the generation of computational analyses.

## Competing statements

Authors declare no competing interests.

## Figure legends

**Figure S1.**
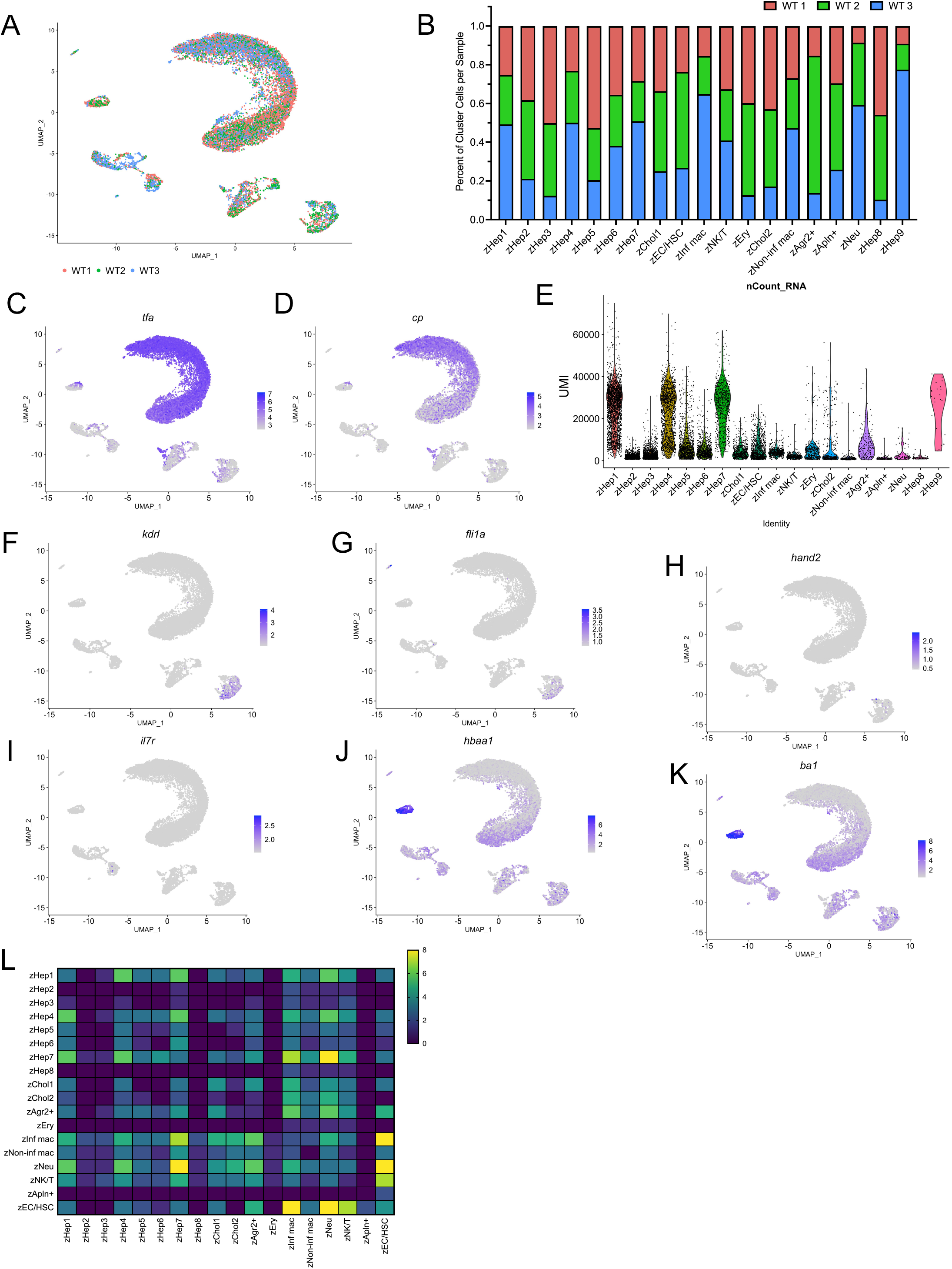
Characterization of the zebrafish adult liver cell clusters. **A**. UMAP of WT adult zebrafish liver atlas with cells color coded by sample identity. **B**. Bar graph showing the percentage of cells contributed to each cluster from each sample. **C-D**. UMAP visualization of hepatocyte marker gene (*tfa* and *cp*) expression. **E**. Violin plot of the number of transcripts detected in each cell grouped by cluster. **F-G**. UMAP visualization of endothelial cell marker gene (*kdrl* and *fli1a*) expression. **H**. UMAP visualization of HSC marker gene (*hand2*) expression. **I**. UMAP visualization of lymphocyte marker gene (*il7r*) expression. **J-K**. UMAP visualization of erythrocyte marker gene (*ba1* and *hbaa1*) expression. **L**. Heatmap of total number of significant receptor-ligand interactions between each cluster as determined by CellPhoneDB analysis performed on the WT zebrafish liver scRNA-seq atlas.

**Figure S2.**
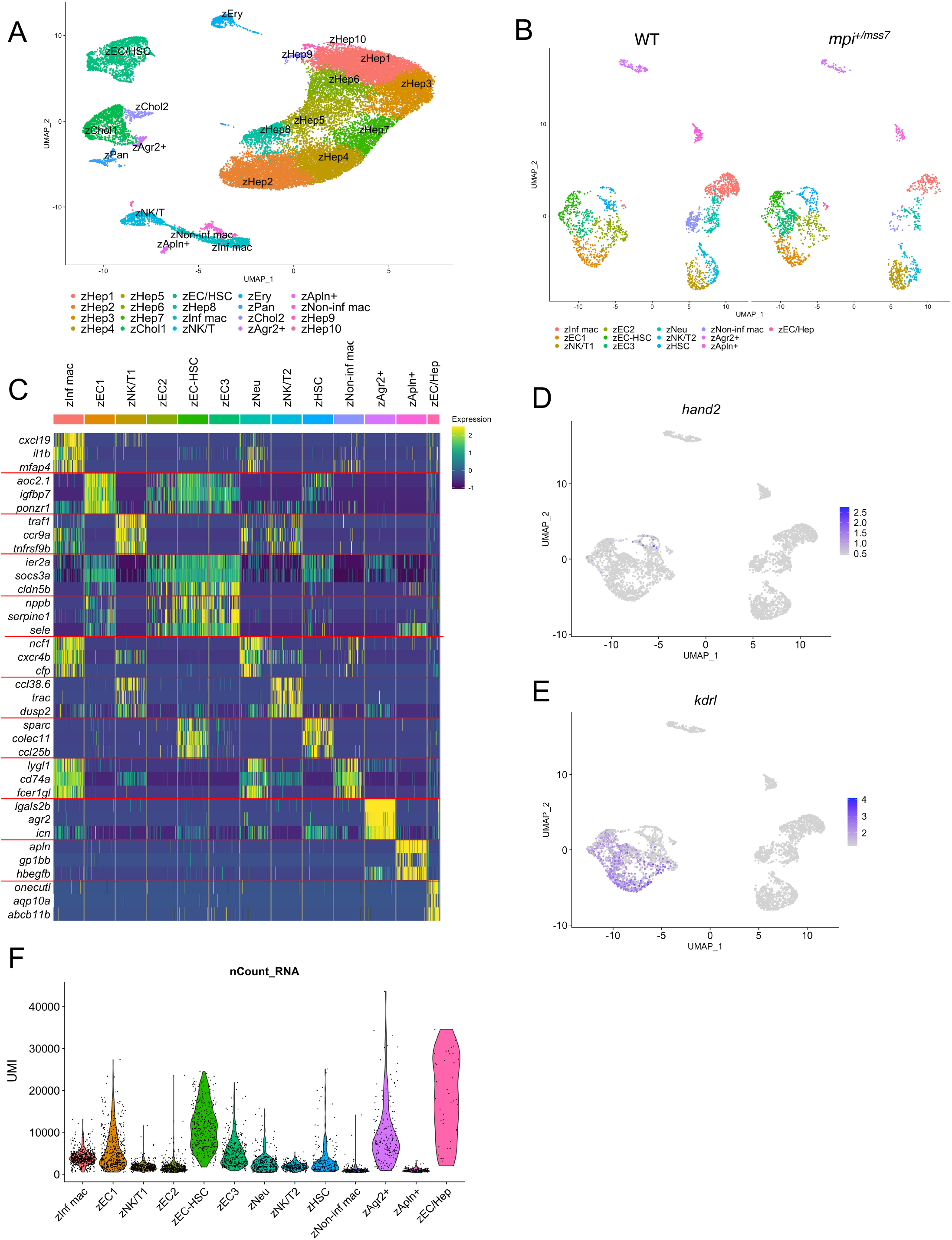
Characterization of WT and *mpi* ^*+/mss7*^ HSC subset of scRNA-seq data. **A**. UMAP of WT and *mpi*^*+/mss7*^ liver cells. **B**. UMAP visualization of 13 clusters of HSCs and their predicted interacting partners, split by genotype. **C**. Heatmap of three of the top differentially expressed genes for each cluster. **D**. UMAP visualization of hepatic stellate cell marker gene (*hand2)* and **E**. endothelial cell marker gene (*kdrl*) expression. **F**. Violin plot of the number of transcripts detected in each cell grouped by cluster.

**Figure S3.**
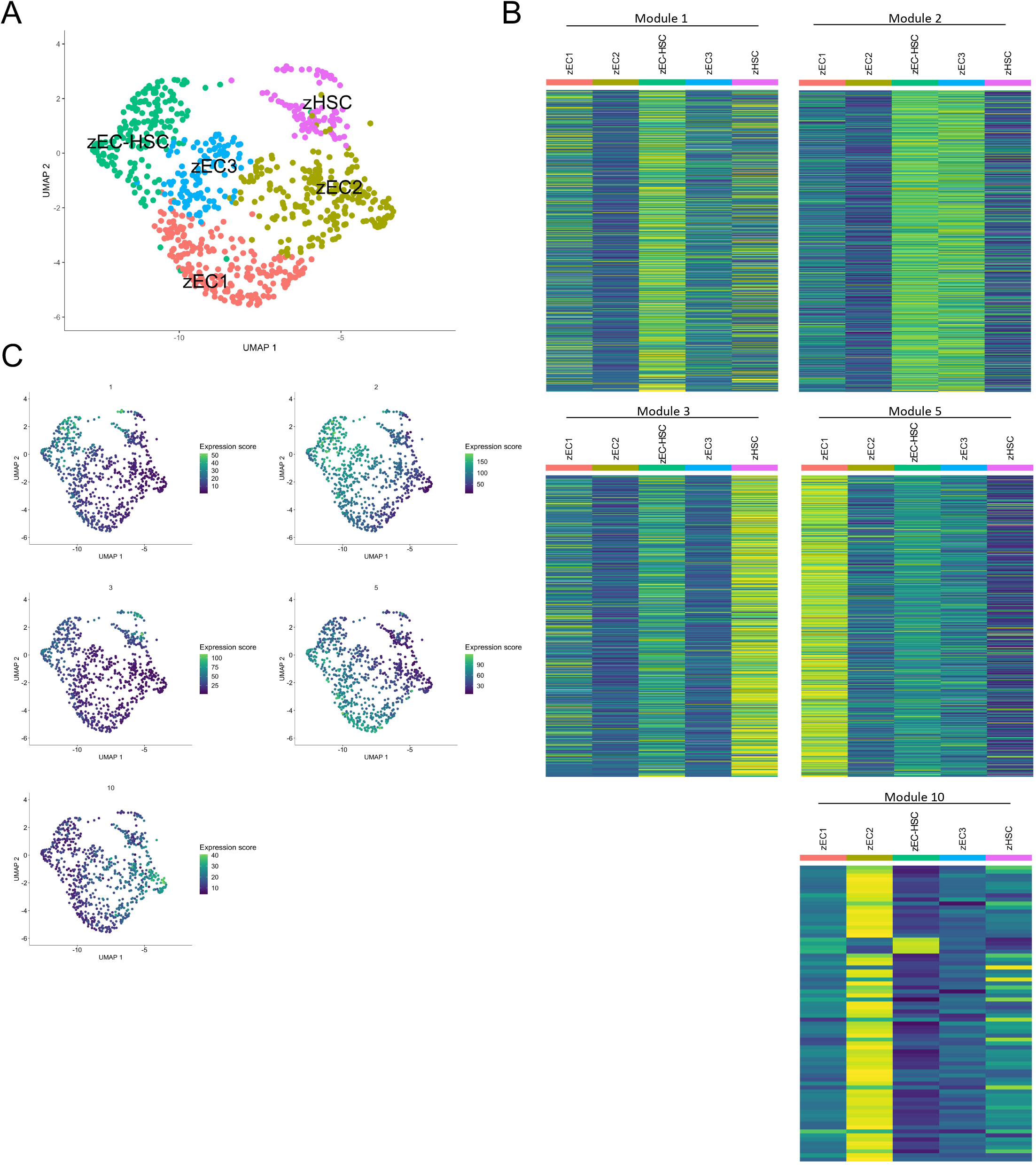
Identifying unique transcriptional profiles for zebrafish HSCs and endothelial cells. **A**. UMAP of WT zebrafish HSC and endothelial cell clusters used for Monocle analysis. **B**. Heatmaps showing expression of the genes that comprise select gene modules that show specificity for a given cluster. **C**. UMAP visualization of the expression of each module across HSC and endothelial cell clusters.

**Figure S4.**
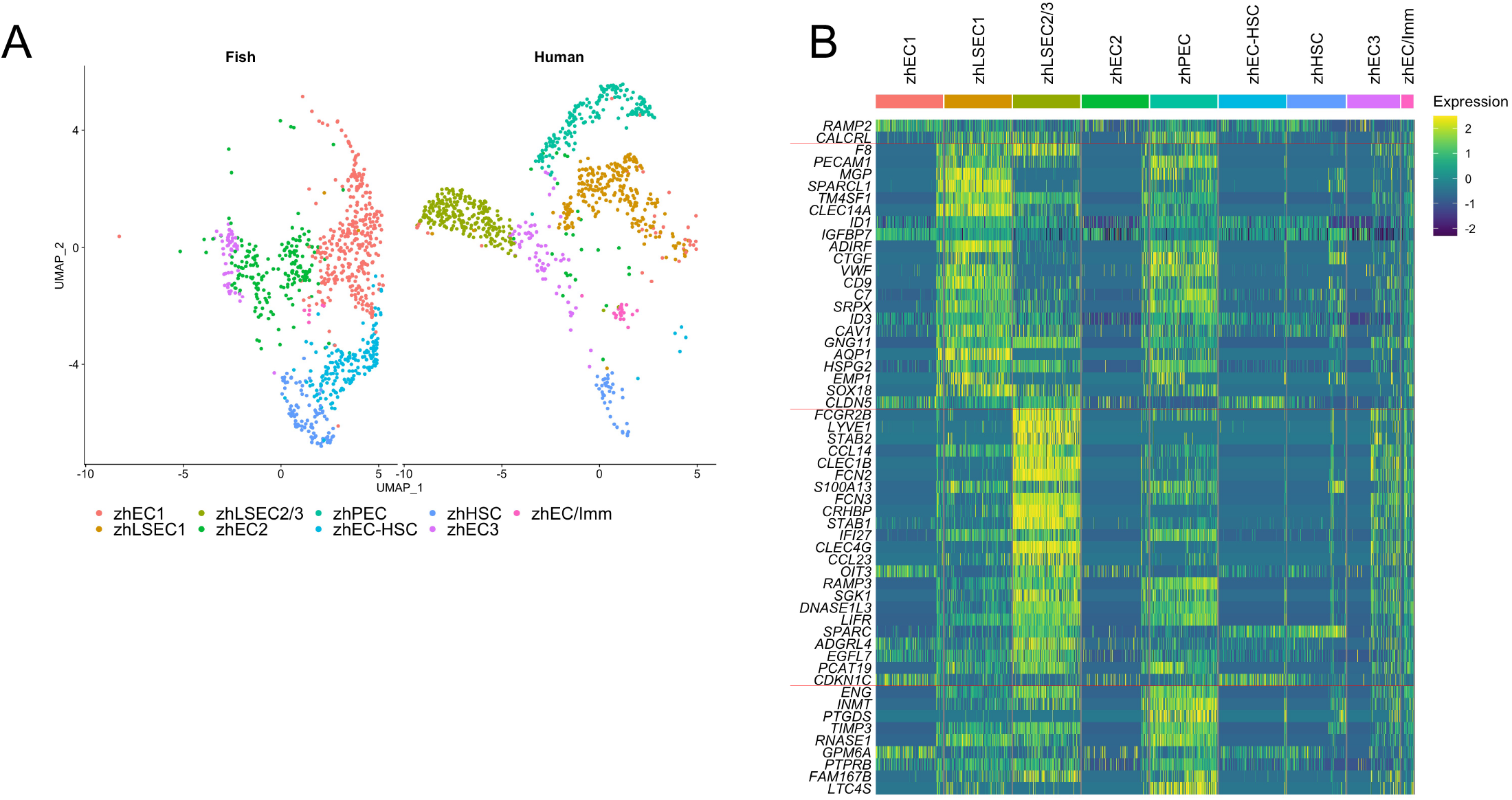
Comparing the transcriptional profiles of human and zebrafish endothelial cells and HSCs. **A**. UMAP visualization of 9 clusters of endothelial cells and HSCs comprised of human and zebrafish cells, split by genotype. **B**. Heatmap of marker genes for human zone 1 LSECs, zone 2/3 LSECs, and portal endothelial cells.

**Figure S5.**
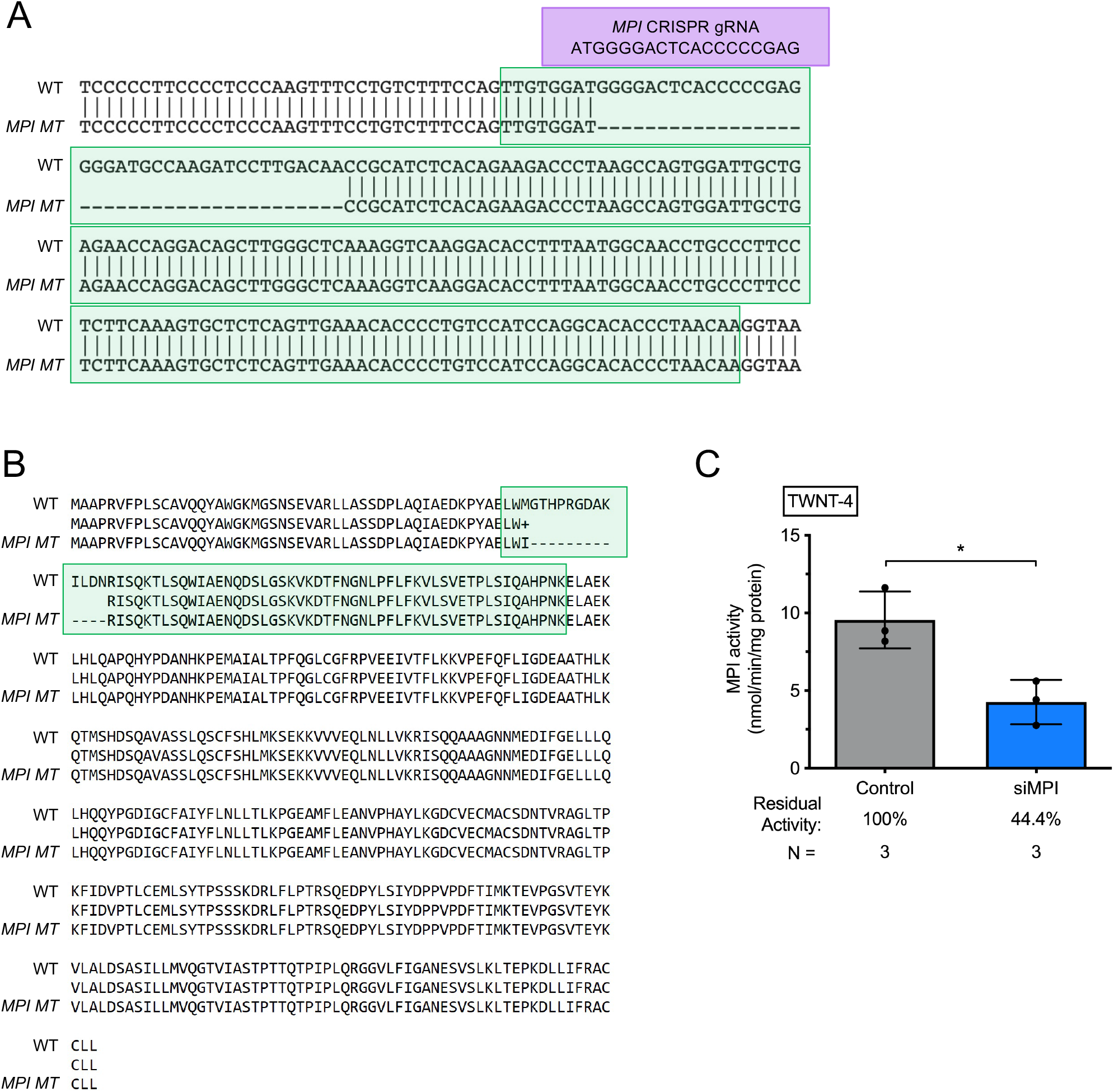
Depleting MPI in human hepatic stellate cell lines. **A**. Alignment of WT *MPI* genomic DNA sequence aligned to the sequence of the mutant *MPI* allele. Exon 3 is highlighted in green and the gRNA target site is in purple. **B**. Alignment of WT *MPI* amino acid sequence aligned to the sequence of the mutant *MPI* allele. Exon 3 is highlighted in green. **C**. Bar graph showing MPI enzymatic activity in control and siMPI TWNT-4s. * p < 0.05

**Table S1**. Marker gene expression for clusters in the WT zebrafish liver atlas.

**Table S2**. Cells per sample in each cluster in the WT zebrafish liver atlas.

**Table S3**. Marker gene expression for clusters in the human and zebrafish liver joint clustering.

**Table S4**. Cells per species in each cluster in the human and zebrafish liver joint clustering.

**Table S5**. Marker gene expression for clusters in the WT and *mpi*^*+/mss7*^ zebrafish liver joint clustering.

**Table S6**. Marker gene expression for clusters in the WT and *mpi*^*+/mss7*^ zebrafish zEC/HSC subset clustering.

**Table S7**. Cells per species in each cluster in the WT and *mpi*^*+/mss7*^ zebrafish zEC/HSC subset clustering.

**Table S8**. Differentially expressed genes in focal regions of WT zebrafish endothelial cells and HSCs in UMAP.

**Table S9**. Modules of gene co-expression for WT zebrafish HSC and endothelial cell clusters.

**Table S10**. Marker gene expression for clusters in the human and WT zebrafish HSC and endothelial cell joint clustering.

**Table S11**. Cells per species in each cluster in the human and WT zebrafish HSC and endothelial cell joint clustering.

**Table S12**. Differentially expressed genes in *mpi*^*+/mss7*^ zebrafish HSCs vs. WT.

**Table S13**. Differentially expressed genes in *MPI MT* LX-2 cells vs. WT.

**Table S14**. Differentially expressed genes in siMPI TWNT-4 cells vs. control.

## References

1. Y. A. Lee, M. C. Wallace, S. L. Friedman, Pathobiology of liver fibrosis: a translational success story. Gut 64, 830–841 (2015).

2. T. Higashi, S. L. Friedman, Y. Hoshida, Hepatic stellate cells as key target in liver fibrosis. Advanced Drug Delivery Reviews 121, 27–42 (2017).

3. J. E. Puche, Y. Saiman, S. L. Friedman, Hepatic Stellate Cells and Liver Fibrosis. Comprehensive Physiology 3, 20 (2013).

4. M. C. H. Janssen, et al., Successful liver transplantation and long-term follow-up in a patient with MPI-CDG. Pediatrics 134, e279–283 (2014).

5. J. Jaeken, et al., Phosphomannose isomerase deficiency: a carbohydrate-deficient glycoprotein syndrome with hepatic-intestinal presentation. Am J Hum Genet 62, 1535– 1539 (1998).

6. P. de Lonlay, N. Seta, The clinical spectrum of phosphomannose isomerase deficiency, with an evaluation of mannose treatment for CDG-Ib. Biochim Biophys Acta 1792, 841–843 (2009).

7. E. Trefts, M. Gannon, D. H. Wasserman, The liver. Curr Biol 27, R1147–R1151 (2017).

8. S. K. Asrani, H. Devarbhavi, J. Eaton, P. S. Kamath, Burden of liver diseases in the world. Journal of Hepatology 70, 151–171 (2019).

9. J. Chu, K. C. Sadler, New school in liver development: lessons from zebrafish. Hepatology 50, 1656–1663 (2009).

10. W. Goessling, K. C. Sadler, Zebrafish: An Important Tool for Liver Disease Research. Gastroenterology 149, 1361–1377 (2015).

11. D.-H. Pham, C. Zhang, C. Yin, Using Zebrafish to Model Liver Diseases-Where Do We Stand? Curr Pathobiol Rep 5, 207–221 (2017).

12. S. de Oliveira, et al., Metformin modulates innate immune-mediated inflammation and early progression of NAFLD-associated hepatocellular carcinoma in zebrafish. Journal of Hepatology 70, 710–721 (2019).

13. O. Tsedensodnom, A. M. Vacaru, D. L. Howarth, C. Yin, K. C. Sadler, Ethanol metabolism and oxidative stress are required for unfolded protein response activation and steatosis in zebrafish with alcoholic liver disease. Dis Model Mech 6, 1213–1226 (2013).

14. T. E. North, et al., PGE2-regulated wnt signaling and N-acetylcysteine are synergistically hepatoprotective in zebrafish acetaminophen injury. Proc Natl Acad Sci U S A 107, 17315– 17320 (2010).

15. N. Shtraizent, et al., MPI depletion enhances O-GlcNAcylation of p53 and suppresses the Warburg effect. eLife 6, e22477 (2017).

16. C. DeRossi, et al., Mannose Phosphate Isomerase and Mannose Regulate Hepatic Stellate Cell Activation and Fibrosis in Zebrafish and Humans. Hepatology 70, 2107–2122 (2019).

17. G. Chen, B. Ning, T. Shi, Single-Cell RNA-Seq Technologies and Related Computational Data Analysis. Front. Genet. 10 (2019).

18. B. Hwang, J. H. Lee, D. Bang, Single-cell RNA sequencing technologies and bioinformatics pipelines. Exp Mol Med 50, 1–14 (2018).

19. N. Aizarani, et al., A Human Liver Cell Atlas reveals Heterogeneity and Epithelial Progenitors. Nature 572, 199–204 (2019).

20. O. Krenkel, J. Hundertmark, T. P. Ritz, R. Weiskirchen, F. Tacke, Single Cell RNA Sequencing Identifies Subsets of Hepatic Stellate Cells and Myofibroblasts in Liver Fibrosis. Cells 8, 503 (2019).

21. S. A. MacParland, et al., Single cell RNA sequencing of human liver reveals distinct intrahepatic macrophage populations. Nature Communications 9, 4383 (2018).

22. V. L. Payen, et al., Single-cell RNA sequencing of human liver reveals hepatic stellate cell heterogeneity. JHEP Reports 3, 100278 (2021).

23. J. Zhao, et al., Single-cell RNA sequencing reveals the heterogeneity of liver-resident immune cells in human. Cell Discov 6, 1–19 (2020).

24. S. Nayar, et al., A myeloid–stromal niche and gp130 rescue in NOD2-driven Crohn’s disease. Nature 593, 275–281 (2021).

25. Y. Hao, et al., Integrated analysis of multimodal single-cell data. Cell 184, 3573-3587.e29 (2021).

26. T. Stuart, et al., Comprehensive Integration of Single-Cell Data. Cell 177, 1888-1902.e21 (2019).

27. A. Butler, P. Hoffman, P. Smibert, E. Papalexi, R. Satija, Integrating single-cell transcriptomic data across different conditions, technologies, and species. Nat Biotechnol 36, 411–420 (2018).

28. R. Satija, J. A. Farrell, D. Gennert, A. F. Schier, A. Regev, Spatial reconstruction of single-cell gene expression data. Nat Biotechnol 33, 495–502 (2015).

29. G. M. Her, C.-C. Chiang, W.-Y. Chen, J.-L. Wu, In vivo studies of liver-type fatty acid binding protein (L-FABP) gene expression in liver of transgenic zebrafish (Danio rerio). FEBS Letters 538, 125–133 (2003).

30. R. Mudbhary, et al., UHRF1 Overexpression Drives DNA Hypomethylation and Hepatocellular Carcinoma. Cancer Cell 25, 196–209 (2014).

31. B. J. Wilkins, M. Pack, Zebrafish Models of Human Liver Development and Disease. Compr Physiol 3, 1213–1230 (2013).

32. S. Korzh, A. Emelyanov, V. Korzh, Developmental analysis of ceruloplasmin gene and liver formation in zebrafish. Mech Dev 103, 137–139 (2001).

33. R. Gebhardt, M. Matz-Soja, Liver zonation: Novel aspects of its regulation and its impact on homeostasis. World J Gastroenterol 20, 8491–8504 (2014).

34. R. Manco, S. Itzkovitz, Liver zonation. Journal of Hepatology 74, 466–468 (2021).

35. I. Manfroid, et al., Zebrafish sox9b is crucial for hepatopancreatic duct development and pancreatic endocrine cell regeneration. Developmental Biology 366, 268–278 (2012).

36. S. Lepreux, P. Bioulac-Sage, E. Chevet, Differential expression of the anterior gradient protein-2 is a conserved feature during morphogenesis and carcinogenesis of the biliary tree. Liver International 31, 322–328 (2011).

37. M. A. Thompson, et al., The cloche and spadetail genes differentially affect hematopoiesis and vasculogenesis. Dev Biol 197, 248–269 (1998).

38. N. D. Lawson, B. M. Weinstein, In Vivo Imaging of Embryonic Vascular Development Using Transgenic Zebrafish. Developmental Biology 248, 307–318 (2002).

39. C. Yin, K. J. Evason, J. J. Maher, D. Y. R. Stainier, The basic helix-loop-helix transcription factor, heart and neural crest derivatives expressed transcript 2, marks hepatic stellate cells in zebrafish: Analysis of stellate cell entry into the developing liver. Hepatology 56, 1958–1970 (2012).

40. J. Rougeot, et al., RNAseq Profiling of Leukocyte Populations in Zebrafish Larvae Reveals a cxcl11 Chemokine Gene as a Marker of Macrophage Polarization During Mycobacterial Infection. Front. Immunol. 10 (2019).

41. D. M. Mitchell, C. Sun, S. S. Hunter, D. D. New, D. L. Stenkamp, Regeneration associated transcriptional signature of retinal microglia and macrophages. Sci Rep 9, 4768 (2019).

42. G. Arango Duque, A. Descoteaux, Macrophage Cytokines: Involvement in Immunity and Infectious Diseases. Front. Immunol. 0 (2014).

43. N. Iwanami, et al., Genetic Evidence for an Evolutionarily Conserved Role of IL-7 Signaling in T Cell Development of Zebrafish. The Journal of Immunology 186, 7060–7066 (2011).

44. J. C. Moore, et al., T Cell Immune Deficiency in zap70 Mutant Zebrafish. Molecular and Cellular Biology 36, 2868–2876.

45. K. Kulkeaw, D. Sugiyama, Zebrafish erythropoiesis and the utility of fish as models of anemia. Stem Cell Res Ther 3, 55 (2012).

46. C. S. McGinnis, et al., MULTI-seq: sample multiplexing for single-cell RNA sequencing using lipid-tagged indices. Nat Methods 16, 619–626 (2019).

47. N. Anzai, et al., Types of nuclear endonuclease activity capable of inducing internucleosomal DNA fragmentation are completely different between human CD34+ cells and their granulocytic descendants. Blood 86, 917–923 (1995).

48. C. S. Helker, et al., Apelin signaling drives vascular endothelial cells toward a pro-angiogenic state. eLife 9, e55589 (2020).

49. M. J. Siemerink, I. Klaassen, C. J. F. Van Noorden, R. O. Schlingemann, Endothelial Tip Cells in Ocular Angiogenesis. J Histochem Cytochem 61, 101–115 (2013).

50. Q. Liu, et al., Genetic targeting of sprouting angiogenesis using Apln-CreER. Nat Commun 6, 6020 (2015).

51. Y. Xie, et al., Key molecular alterations in endothelial cells in human glioblastoma uncovered through single-cell RNA sequencing. JCI Insight (2021) https://doi.org/10.1172/jci.insight.150861 (August 2, 2021).

52. H. Yokomori, M. Oda, K. Yoshimura, T. Hibi, Enhanced expressions of apelin on proliferative hepatic arterial capillaries in human cirrhotic liver. Hepatology Research 42, 508–514 (2012).

53. T. Tsuchida, S. L. Friedman, Mechanisms of hepatic stellate cell activation. Nat Rev Gastroenterol Hepatol 14, 397–411 (2017).

54. L. D. DeLeve, Liver Sinusoidal Endothelial Cells in Hepatic Fibrosis. Hepatology 61, 1740– 1746 (2015).

55. M. Efremova, M. Vento-Tormo, S. A. Teichmann, R. Vento-Tormo, CellPhoneDB: inferring cell–cell communication from combined expression of multi-subunit ligand–receptor complexes. Nat Protoc 15, 1484–1506 (2020).

56. D. Semela, et al., PDGF signaling through ephrin-B2 regulates hepatic vascular structure and function. Gastroenterology 135, 671–679 (2008).

57. C. Trapnell, et al., The dynamics and regulators of cell fate decisions are revealed by pseudotemporal ordering of single cells. Nat Biotechnol 32, 381–386 (2014).

58. X. Qiu, et al., Reversed graph embedding resolves complex single-cell trajectories. Nat Methods 14, 979–982 (2017).

59. J. Cao, et al., The single-cell transcriptional landscape of mammalian organogenesis. Nature 566, 496–502 (2019).

60. L. McInnes, J. Healy, J. Melville, UMAP: Uniform Manifold Approximation and Projection for Dimension Reduction. 1802.03426 [cs, stat] (2020) (July 19, 2021).

61. V. A. Traag, L. Waltman, N. J. van Eck, From Louvain to Leiden: guaranteeing well-connected communities. Sci Rep 9, 5233 (2019).

62. J. H. Levine, et al., Data-Driven Phenotypic Dissection of AML Reveals Progenitor-like Cells that Correlate with Prognosis. Cell 162, 184–197 (2015).

63. W. Yang, et al., Single-cell transcriptomic analysis reveals a hepatic stellate cell-activation roadmap and myofibroblast origin during liver fibrosis. Hepatology n/a.

64. S. Morini, et al., GFAP expression in the liver as an early marker of stellate cells activation. Ital J Anat Embryol 110, 193–207 (2005).

65. B. Zhang, C.-G. Zhang, L.-H. Ji, G. Zhao, Z.-Y. Wu, Estrogen receptor β selective agonist ameliorates liver cirrhosis in rats by inhibiting the activation and proliferation of hepatic stellate cells. J Gastroenterol Hepatol 33, 747–755 (2018).

66. L. Shan, et al., mTOR Overactivation in Mesenchymal cells Aggravates CCl 4 - Induced liver Fibrosis. Sci Rep 6, 36037 (2016).

67. G. Svegliati-Baroni, et al., Insulin and insulin-like growth factor-1 stimulate proliferation and type I collagen accumulation by human hepatic stellate cells: differential effects on signal transduction pathways. Hepatology 29, 1743–1751 (1999).

68. R. G. Gieling, K. Wallace, Y.-P. Han, Interleukin-1 participates in the progression from liver injury to fibrosis. Am J Physiol Gastrointest Liver Physiol 296, G1324–G1331 (2009).

69. I. Fabregat, et al., TGF-β signalling and liver disease. The FEBS Journal 283, 2219–2232 (2016).

70. Y. Osawa, et al., Tumor Necrosis Factor-α Promotes Cholestasis-Induced Liver Fibrosis in the Mouse through Tissue Inhibitor of Metalloproteinase-1 Production in Hepatic Stellate Cells. PLOS ONE 8, e65251 (2013).

71. D. Liang, et al., Inhibition of EGFR attenuates fibrosis and stellate cell activation in diet-induced model of nonalcoholic fatty liver disease. Biochimica et Biophysica Acta (BBA) - Molecular Basis of Disease 1864, 133–142 (2018).

72. C. Yang, et al., Liver fibrosis: Insights into migration of hepatic stellate cells in response to extracellular matrix and growth factors. Gastroenterology 124, 147–159 (2003).

73. P. Kocabayoglu, et al., β-PDGF receptor expressed by hepatic stellate cells regulates fibrosis in murine liver injury, but not carcinogenesis. J Hepatol 63, 141–147 (2015).

74. J. Fan, et al., Bone morphogenetic protein 4 mediates bile duct ligation induced liver fibrosis through activation of Smad1 and ERK1/2 in rat hepatic stellate cells. Journal of Cellular Physiology 207, 499–505 (2006).

75. C. Yu, et al., Role of Fibroblast Growth Factor Type 1 and 2 in Carbon Tetrachloride-Induced Hepatic Injury and Fibrogenesis. The American Journal of Pathology 163, 1653– 1662 (2003).

76. L. Yang, et al., Vascular endothelial growth factor promotes fibrosis resolution and repair in mice. Gastroenterology 146, 1339-1350.e1 (2014).

77. J. Luo, et al., Vascular endothelial growth factor promotes the activation of hepatic stellate cells in chronic schistosomiasis. Immunol Cell Biol 95, 399–407 (2017).

78. M. Sun, T. Kisseleva, Reversibility of liver fibrosis. Clin Res Hepatol Gastroenterol 39 Suppl 1, S60–63 (2015).

79. N. Roehlen, E. Crouchet, T. F. Baumert, Liver Fibrosis: Mechanistic Concepts and Therapeutic Perspectives. Cells 9, 875 (2020).

80. M. L. Richter, et al., Single-nucleus RNA-seq2 reveals functional crosstalk between liver zonation and ploidy. Nat Commun 12, 4264 (2021).

81. Q. Yang, C. Yan, Z. Gong, Interaction of hepatic stellate cells with neutrophils and macrophages in the liver following oncogenic kras activation in transgenic zebrafish. Sci Rep 8, 8495 (2018).

82. C. DeRossi, et al., Ablation of Mouse Phosphomannose Isomerase (Mpi) Causes Mannose 6-Phosphate Accumulation, Toxicity, and Embryonic Lethality *. Journal of Biological Chemistry 281, 5916–5927 (2006).

83. R. J. Willms, J. C. Hocking, E. Foley, A Cell Atlas of Microbe-Responsive Processes in the Zebrafish Intestine. bioRxiv, 2020.11.06.371609 (2020).

84. E. Eden, R. Navon, I. Steinfeld, D. Lipson, Z. Yakhini, GOrilla: a tool for discovery and visualization of enriched GO terms in ranked gene lists. BMC Bioinformatics 10, 48 (2009).

85. F. Supek, M. BoŠnjak, N. Škunca, T. Šmuc, REVIGO Summarizes and Visualizes Long Lists of Gene Ontology Terms. PLOS ONE 6, e21800 (2011).

86. L. Xu, et al., Human hepatic stellate cell lines, LX-1 and LX-2: new tools for analysis of hepatic fibrosis. Gut 54, 142–151 (2005).

87. N. Shibata, et al., Establishment of an Immortalized Human Hepatic Stellate Cell Line to Develop Antifibrotic Therapies. Cell Transplant 12, 499–507 (2003).

88. J. Chu, et al., A zebrafish model of congenital disorders of glycosylation with phosphomannose isomerase deficiency reveals an early opportunity for corrective mannose supplementation. Dis Model Mech 6, 95–105 (2013).

89. M. I. Love, W. Huber, S. Anders, Moderated estimation of fold change and dispersion for RNA-seq data with DESeq2. Genome Biology 15, 550 (2014).

90. R. C. Gentleman, et al., Bioconductor: open software development for computational biology and bioinformatics. Genome Biol 5, R80 (2004).

91. A. Krämer, J. Green, J. Pollard Jr, S. Tugendreich, Causal analysis approaches in Ingenuity Pathway Analysis. Bioinformatics 30, 523–530 (2014).

